# Mapping Responsive Genomic Elements to Heat Stress in a Maize Diversity Panel

**DOI:** 10.1101/2022.06.23.497238

**Authors:** Zhikai Liang, Zachary A. Myers, Dominic Petrella, Julia Engelhorn, Thomas Hartwig, Nathan M. Springer

## Abstract

Many plant species exhibit genetic variation for tolerating environmental stress. A transcriptome investigation of over 100 maize inbreds revealed many *cis*- and *trans*-acting eQTLs that influence the expression response to heat stress. The *cis*-acting eQTL in response to heat stress are identified in genes with differential responses to heat stress between genotypes as well as genes that are only expressed under heat stress. The *cis*-acting variants for heat stress responsive expression likely result from distinct promoter activities and the differential heat responses of the alleles were confirmed for selected genes using transient expression assays. Global foot-printing of transcription factor binding was performed in control and heat stress conditions to document regions with heat-enriched transcription factor binding occupancies. Footprints enriched near proximal regions of characterized heat-responsive genes in a large association panel can be utilized for prioritizing functional genomic regions that regulate genotype-specific responses under heat stress.

## Introduction

Changes in gene expression are one mechanism in plants for tolerating changing environmental conditions ^1^. Heat is an abiotic stress that can affect plants in many ways. Minor changes in average temperatures can result in lower average yield ^2^. Severe heat stress events during reproductive periods can result in major changes in the seed set ^3,4^. Heat stress events at earlier vegetative stages can result in leaf senescence and reduced growth rate ^5,6^.

Transcriptome profiling in plants has identified many genes that exhibit altered transcript abundance in response to abiotic stress ^7–11^. The heat-shock factor (HSF) transcription factors (TFs) are activated in response to heat stress ^12^ and can regulate the expression level of many genes through interactions with *cis*-acting elements. Prior studies have found evidence for enrichment of the HSF binding site motif at up-regulated genes in Arabidopsis ^13^, maize ^14^ and rice ^15^. Some HSFs are stably expressed and become activated in response to heat stress, while many other members of this family are themselves transcriptionally activated in response to heat stress ^12,16,17^. Several other transcription factors and *cis*-regulatory elements have also been implicated in the response to heat stress ^18^. The prior knowledge of potential mechanisms that create altered gene expression in response to heat stress makes this a good system to study variable responses to heat stress.

One difficulty in elucidating the mechanisms of gene expression regulation has been our ability to identify and annotate *cis*-regulatory elements that influence gene expression patterns and levels. In some cases, researchers simply utilize all regions upstream of the transcriptional start site (typically 1-2kb) to search for potential motifs or regulatory elements. Comparisons among related species are also utilized to identify conserved non-coding sequences (CNSs) that may be important for regulation ^19,20^. Recently, many groups have focused on using chromatin properties to identify putative *cis*-regulatory elements ^21–24^. Specifically, regions of accessible chromatin or particular histone modifications can be useful in documenting functionally important *cis*-regulatory elements ^25^. However, for gene expression responses to abiotic stress, it is unclear what proportion of *cis*-regulatory elements are premarked (accessible prior to the stress) or only exhibit altered accessibility following the stress exposure.

Local adaptation in wild species and breeding to improve the resilience of crop plants likely utilize natural variation in gene expression responses to abiotic stresses ^26^. This variation can be attributed to varying activity of *trans*-acting factors such as transcription factors or due to *cis*-acting regulatory variation that can alter how specific genes are regulated. Our understanding of the evolutionary sources of variable *cis*-regulatory elements is limited. The mechanisms by which genes gain, or lose, responsiveness to an environmental stimulus is unclear. It is assumed that a large portion of *cis*-regulatory informtion is provided through transcription factor binding sites (TFBSs). However, it is not understood how novel TFBSs are created, or lost, at specific genes. Single nucleotide polymorphisms (SNPs) might contribute to altered *cis*-regulatory features, but it is also possible that insertion / deletion (InDel) polymorphisms, including transposable element (TE) insertions, play a major role in changing the composition of *cis*-regulatory elements. Several studies have compared the gene expression responses to heat stress in different genotypes ^8,14,27–29^. While many genes exhibit similar responses to heat stress in different genotypes there are also examples of genes that exhibit significant allelic variation for responses ^8,14^. The use of hybrid genotypes and allele-specific expression analyses has found evidence for both *cis*- and *trans*-regulatory variation. A comparison of the alleles suggests that structural variation may play important roles in driving variable responsiveness but this has not been investigated in detail.

In this study, we utilized a panel of 102 resequenced maize genotypes grown in control and heat stress conditions to study variation of gene expression and associated regulatory genomic elements in response to heat. Both genomic and transcriptomic data were utilized to document eQTLs and their regulated genes. In particular, *cis*-eQTL that influence variable responsiveness to heat stress were identified to document allelic variation for heat induced expression. This approach was demonstrated as an efficient way to document hundreds of *cis*-allelic variations for heat stress response. The analysis of *cis*-regulatory variation for heat responses using eQTL was complemented with a genome-wide transcription factor footprinting assay in B73. MNase-defined cistrome-Occupancy Analysis (MOA-seq) data ^24^ was generated in both control and heat stress conditions to document regions with altered chromatin accessibility and transcription factor occupancy. We find that many heat-responsive genes exhibit changes in chromatin accessibility and transcription factor occupancy in the heat stress condition relative to control. The combination of these two approaches provided opportunities to identify high quality candidate causal regions that could explain the allelic variation for heat responsiveness.

## Results

### Genome-wide markers associated with photosynthetic parameters

A subpanel of 102 diverse maize genotypes including stiff stalk, non-stiff stalk and iodent sub-populations representing the Wisconsin Diversity Panel (WiDiv) in maize ^30,31^ were selected for characterizing genotype responses to heat stress (Figure 1a). A set of 1,132,322 variants (1,032,834 SNPs + 99,488 InDels) was employed for this population (see methods for filtering details). The first five principal components (PCs) calculated from the genotype matrix could explain 17.1% of population structure variation in the selected 102 genotypes. Seedlings (14 days after sowing) for each of these genotypes were subjected to a 4 hour heat stress at 40C. The third leaf for each plant was also assessed for three chlorophyll fluorescence-based parameters including 1) light-adapted effective quantum yield of PSII [Y(II)], and two non-photochemical dissipation routes, 2) non-regulated energy dissipation at PSII [Y(NO)], and 3) regulated energy dissipation at PSII [Y(NPQ)] in control and heat stress conditions. Higher Y(II) values indicated less photosynthetic stress, high Y(NPQ) values indicated ongoing photochemical stress that was being mitigated through carotenoid-based energy dissipation, and high Y(NO) values indicated ongoing photochemical stress that was not being mitigated. On average, plants exhibited lower Y(NO) values and higher Y(II) values under the heat stress condition compared to the control condition, while the center of recorded Y(NPQ) did not show a significant shift between control and heat conditions (Figure 1b; Figure S1). All three traits [Y(II), Y(NPQ), and Y(NO)] exhibited higher broad-sense heritability (H^2^) in control (0.692, 0.682 and 0.716) compared to heat stressed plants (0.538, 0.563 and 0.642), indicating more variation for these traits in heat stress condition compared to control plants.

**Figure 1.**
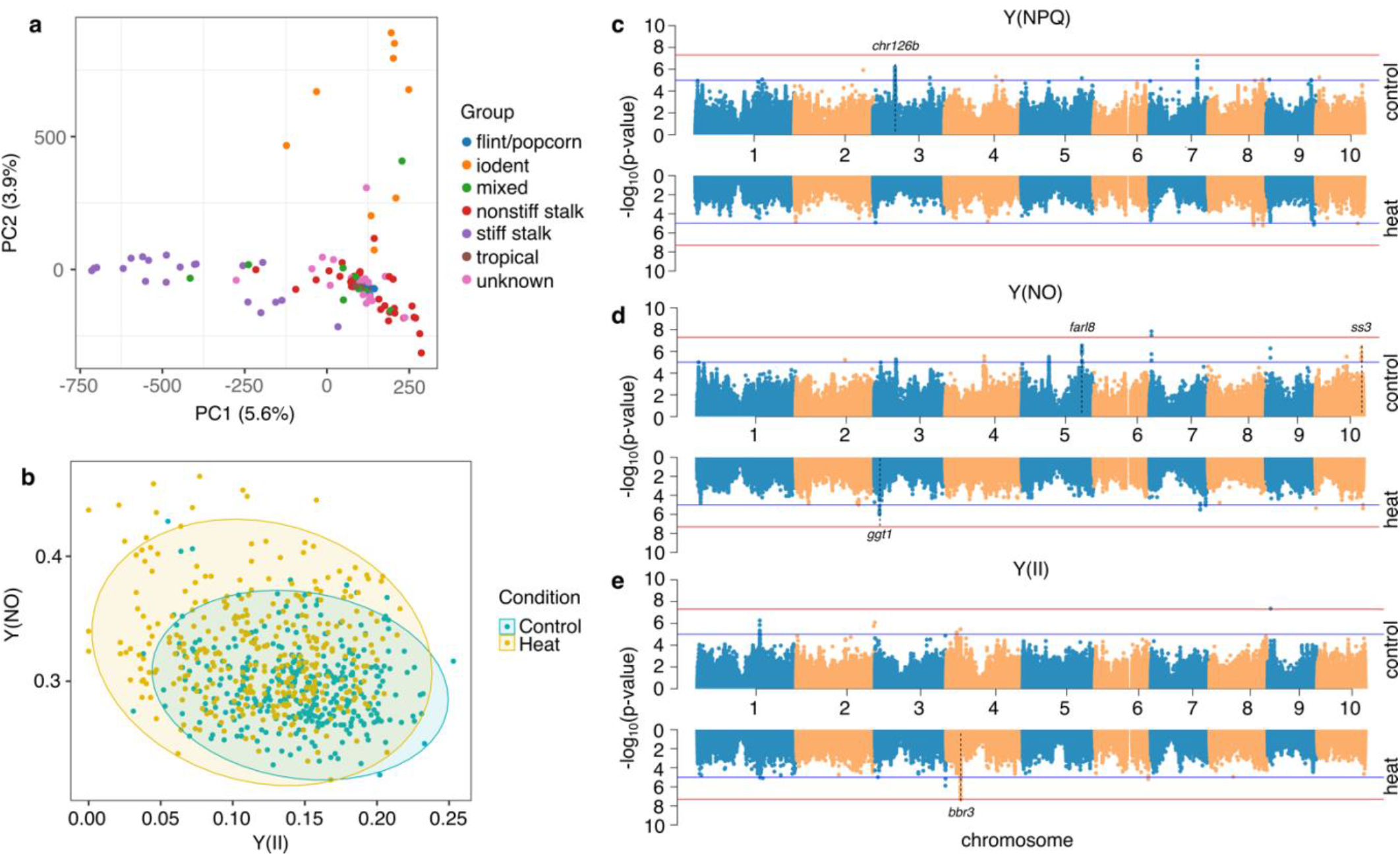
Phenotypic responses to heat stress within a maize diversity panel. (a) Genotypic variants segregated 102 genotypes employed in this study using principal component analysis; (b) Plants showed a distinct pattern using two photosynthetic parameters -- Y(NO) and Y(II) -- measured under control and heat conditions; (c) Associated loci were detected under control and heat conditions using GWAS for Y(NPQ). Two thresholds including 2.2e^−6^ (blue) and 1.1e^−7^ (red) were applied for detecting significantly associated loci; (d) Same as (c) for Y(NO); (e) Same as (c) for Y(II).

The BLUP (Best Linear Unbiased Predictor) was applied to represent the phenotypic value of each genotype per trait in each condition in order to remove underlying variation. BLUPs from the chlorophyll fluorescence-based traits (Supplemental Table S1) were used to perform GWAS (Genome-Wide Association Study) using the same set of >1M variants in this population. We identified candidate association loci under two levels of thresholds (2.2e^−6^ and 1.1e^−7^, see Methods) using the FarmCPU model ^32^. Distinct patterns of GWAS significant hits were observed for traits taken from control and heat stress conditions (Figure 1c-e). In total, 20 candidate genes were identified to be associated with at least one of the significant loci (Supplemental Table S2). Several interesting candidate hits included SNPs located within introns of the *farl8* (FAR1-like transcription factor 8, *Zm00001d017164*) gene that were associated with Y(NO) under the control condition. Orthologs of *Zm00001d017164* in *Arabidopsis* -- *AT1G52520* and *AT1G80010* were demonstrated to show visible phenotypes under abiotic stress ^33^. A SNP located ~1kb downstream of the transcription factor -- *bbr3* (BBR/BPC-transcription factor 3, *Zm00001d049800*) -- was uniquely identified to be significantly associated with Y(II) measured under the heat condition. *Zm00001d049800* encoded as a member of the BASIC PENTACYSTEINE (BPC) proteins and its ortholog *AT1G14685* in *Arabidopsis* was found to be involved in the salt stress response regulation ^34^.

### Transcriptome responses to heat stress in a subset of WiDiv panel

We proceeded to evaluate the transcriptional response to a heat stress event in the same panel of genotypes. Seedlings (14 days after sowing) were subjected to control conditions or 4 hours of heat stress (40C) and leaf tissue collected from a pool of 2-3 individuals for each treatment / genotype combination was used for RNA-seq. The RNA-seq data was aligned to the maize B73 AGPv4 reference genome and all samples exhibited high overall alignment rates, ranging from 84.5% to 98.6% (Supplemental Table S3). After removing genes with no read counts in any samples, we obtained a gene count matrix with reads assigned to a set of 39,511 maize AGPv4 gene models. In order to assess the consistency between samples during the collection, three biological replicates of B73 were separately collected at early, middle and late collection time points in each condition. The average pairwise spearman correlations between replicates was 0.987 and 0.972 in control and heat respectively (Figure S2), indicating a high repeatability of samples during collections. The overall variability in the transcriptomes of all samples were assessed using a principal component analysis (Figure 2a). The first principal component separated samples based on treatment, suggesting a significant impact from the heat stress (Figure 2a). The second principal component provided a modest separation based on the known population structure (Figure 2a). We hypothesized genotypes with less transcriptome shift (based on principal component 1) in response to heat stress might exhibit a more similar distribution of BLUPs for photosynthetic parameters between control and heat condition. A comparison of the absolute difference of the first principal component values between control and heat condition based on transcriptome data and the correlation coefficient of the BLUP values for the photosynthetic traits revealed a significantly positive correlation, suggesting an overall positive correlation between phenotypic and transcriptomic responses (Figure 2b; Figure S3). A GO (Gene Ontology) enrichment analysis on genes with significant up-regulation in response to heat in B73 revealed many significant GO enrichments that were associated with heat responsive functions (i.e. GO:0009408 “response to heat”; GO:0010286 “heat acclimation”; GO:0031072 “heat shock protein binding”) (Supplemental Table S4), providing evidence of a strong heat stress response in the B73 samples.

**Figure 2.**
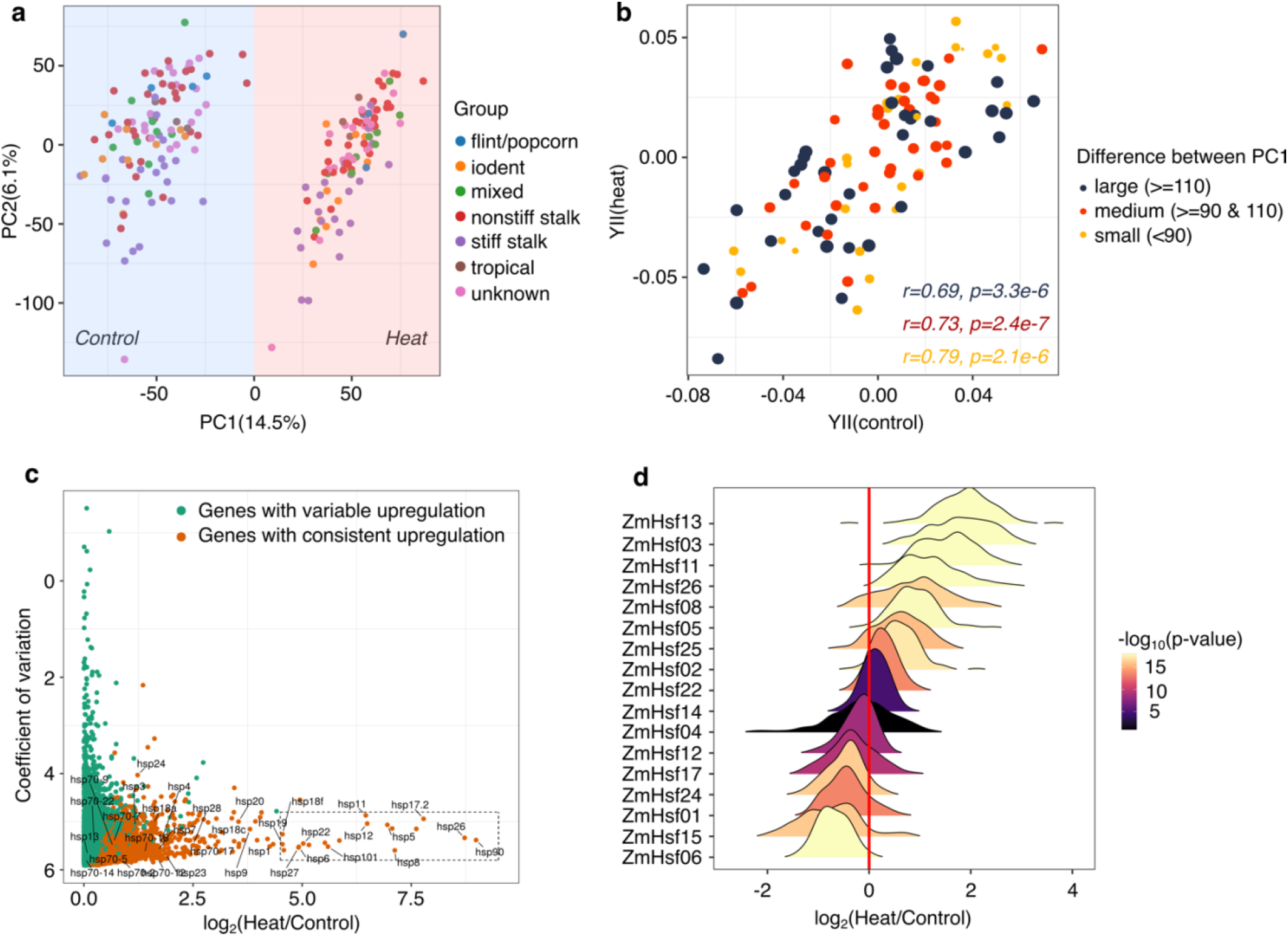
Transcriptomic diversity of studied maize genotypes between heat and control conditions. (a) The PCA analysis separated expressions of identical genotypes in control and heat conditions using normalized read counts of expressed genes; (b) Genotypes with variable levels of absolute PC1 differences were split into three classes: large, medium and small. The correlation of BLUPs generated from Y(II) in control and heat condition was separately calculated; (c) Across 102 studied genotypes, we identified upregulated genes based on the fold change of median expression value between heat and control. We then visualized these upregulated genes based on their coefficient of variations and log_2_ fold change of heat to control. Heat shock proteins were labeled in this graph and the blocking area demonstrated a clustered region with heat shock proteins; (d) Distributions of log_2_ fold changes of heat-to-control normalized expression values for heat shock factors identified in this study.

The analysis of the full dataset revealed 2,628 genes that exhibited consistent up-regulation in response to heat in the majority of genotypes (see Methods for details). A comparison of the mean response to heat stress and the coefficient of variation (CV) for expression level in heat stress for the 2,628 genes highlighted a subset of genes with strong expression response and limited variability in the response for different genotypes (Figure 2c). Many of the genes with the highest expression response and relatively low variation among genotypes were previously identified as heat shock proteins (hsps). There were 40 HSPs identified in the maizeGDB database (https://www.maizegdb.org) and 25 of these HSPs were detected to be consistently upregulated with high expression values and low CVs under the heat stress (Figure 2c). Six additional genes (*Zm00001d004243, Zm00001d022630, Zm00001d031436, Zm00001d033990, Zm00001d039933* and *Zm00001d048592*) that were not annotated as hsps, but also exhibited strong and consistent responses to heat stress and might play important roles in response to heat stress. The heat shock factors (HSFs) were a class of transcription factors that were known to play important roles in gene expression responses to heat stress and activated HSPs. Of 31 previously reported non-redundant HSFs in maize ^17^, 17 of them exhibited detectable expression in our dataset. Eight of these HSFs (*ZmHsf02, ZmHsf25, ZmHsf05, ZmHsf08, ZmHsf26, ZmHsf11, ZmHsf03* and *ZmHsf13*) were consistently upregulated in response to heat stress across a hundred of genotypes (Figure 2d). Interestingly, *ZmHsf06* was downregulated across all genotypes used in this study and also exhibited reduced expression in response to heat stress at multiple timepoints in B73 in a previous study ^35^ (Figure S4). The results suggested that the heat-stress event resulted in a robust response to heat stress in this diverse panel and provided an opportunity to further examine variable responses in more diverse genotypes.

### Characterization of variable gene expression responses to heat stress

To map regulatory variants that might influence gene expression levels or responses to heat stress in this panel, we retained genes with counts per million (CPM) > 1 in at least 10% genotypes, resulting in 20,255 and 20,306 genes expressed in control and heat conditions, respectively. Of these genes, 19,642 genes were commonly expressed in both conditions, while 613 and 664 genes were uniquely detected as expressed in control or heat conditions. Of 664 genes uniquely detected in heat condition, three were heat shock factors (*ZmHsf03, ZmHsf11* and *ZmHsf26*). We performed eQTL mapping separately using control or heat expression data to map the genomic elements influencing transcript levels of expressed genes. Significantly associated SNPs located within 1Mb distance from the targeted gene were classified as *cis*-eQTL, while SNPs located >1Mb from the target gene were classified as *trans-eQTL*. Potential *trans-eQTL* hotspots were assessed by counting the number of significant *trans*-interactions for each 10kb window in the genome (Figure S5). Several chromosomal regions with at least 10 *trans*-interactions were commonly detected for both heat and control conditions. Three 10kb regions on chromosomal 9 were detected to be interacted with gene expression distantly under heat stress. However, the detected ‘hotspots’ had somewhat limited numbers of targets (<20) and the power to detect *trans*-eQTL hotspots in this study might be limited by our sample size. Consistent with prior studies ^36^, identified *cis*-eQTLs in this study could explain more variance of gene expression than identified *trans*-eQTLs in both control and heat conditions (Figure S6). The *cis*-eQTL analysis identified 10,548 and 10,391 significant *cis*-eQTL regulated genes (eGenes) for control and heat condition, respectively. The *cis*-eQTLs detected for genes that were only expressed in heat stress conditions were particularly interesting, as they reflected variable levels of gene expression activation under heat stress. There were 518 of 664 <u>h</u>eat <u>e</u>xpressed <u>o</u>nly (heo) genes that had signifcinat *cis*-eQTL (heo-*cis*-eQTL) and we subsequently referred to these as heo-eGenes (Supplemental Table S5) (Figure S7). The heo-eGenes included a number of transcription factors, such as *zhd20, iaa37, abi33, gata17, sbp18, wrky35, bhlh110, dof27, ereb196*. In addition, two heat shock factors -- *ZmHsf03* and *ZmHsf26* -- were detected as heo-eGenes.

To identify *cis*-regulatory variation associated with heat responsiveness per gene in 19,642 commonly expressed genes under both control and heat conditions, a linear mixed model was incorporated with covariates to detect the interaction effect between each pair of *cis*-eQTL and eGene. To avoid redundancy testing, only the most signficant *cis*-eQTLs in either condition (control or heat) were retained for each gene (two different *cis*-eQTLs might be selected for testing if they had equivalently significant p-values; see Methods), resulting in 15,588 tested gene/*cis*-eQTL pairs. Of tested *cis*-eQTLs, the majority had positive correlations for effects between control and heat conditions (Figure 3a). However, a set of 273 *cis*-eQTLs with significant differences in effects between control and heat conditions were defined as responsive eQTLs (reQTLs) (Figure 3a). These reQTLs provided examples of genes with significant differences in response to heat stress due to *cis*-regulatory variation for responsiveness. Compared to all *cis*-eQTLs identified in either of conditions or selected top significant *cis*-eQTLs for reQTLs identification, reQTLs *per se* were more likely to be distributed around TSS (Transcriptional Start Site) regions and much less likely to be located >3Kb downstream of regulated genes (Figure 3b).

**Figure 3.**
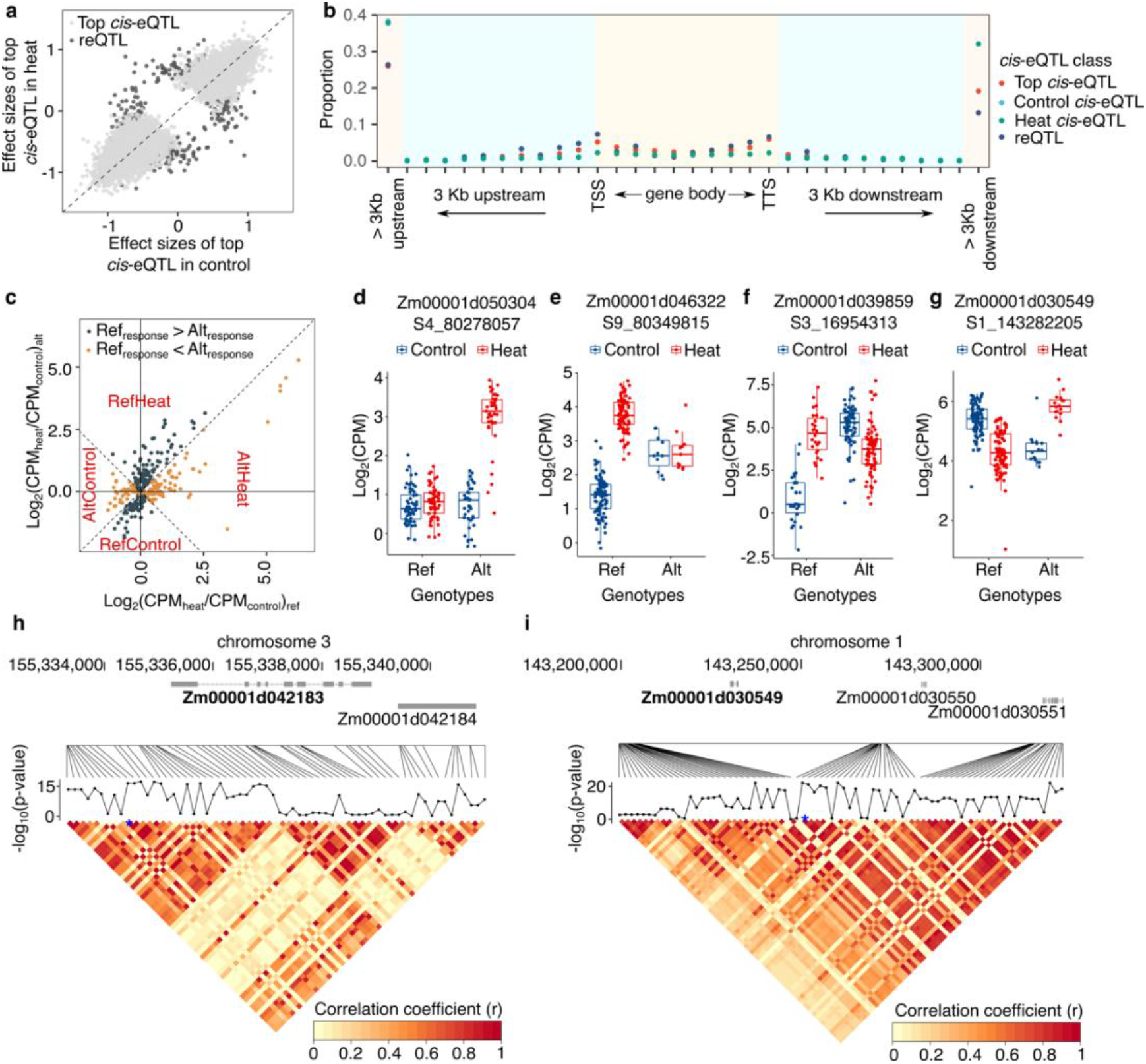
Detected responsive genes and eQTLs in the studied maize diversity panel. (a) Correlation of effect sizes of selected top significant *cis*-eQTLs versus identified responsive eQTLs (reQTLs); (b) Distribution of *cis*-eQTLs nearby annotated genes. Control *cis*-eQTLs represented *cis*-eQTLs identified using control expression data and heat *cis*-eQTLs represented *cis*-eQTLs identified using heat expression data. The y-axis indicated the proportion of each *cis*-eQTL class. TSS: Transcription Start Sites; TTS: Transcription Terminal Sites; (c) Log_2_ fold changes between heat and control for identified responsive genes (reGenes). The x-axis indicated log_2_ fold changes of expressions of genes carrying reference allele between heat and control. The y-axis indicated log_2_ fold changes of expressions of gene carrying alternative alleles between heat and control. The CPM of 1 was added to both denominator and numerator to the ratio to enable the calculation. Based on the ratio of median expression in heat to control, reGenes were subclassified into RefControl, RefHeat, AltControl and AltHeat; (d-g) Four cases shew different responsive genes; (h) Connected curve indicated the significant p-value of interacted term between SNPs and targeted reGene (bold). The heatmap showed the pairwise correlation of SNPs in the displayed genomic region. Blue star indicated the top selected *cis*-eQTL for genome-wide reQTL identification analysis; (i) Same as the panel h, but showing gene *Zm00001d030549*.

The reQTL regulated genes (reGenes) were further assessed and classified into two categories based on the strength of responsiveness under heat stress: reference alleles with greater response or alternative allele with greater response to heat stress. Both of these groups could be further subdivided based on whether the more responsive allele was up- or down-regulated in response to heat stress. Of total 258 identified reGenes (Supplemental Table S6), 54.4% of these reGenes were examples with reference alleles that have consistently higher response to heat compared to the alternate allele while the other 45.6% exhibited consistently higher response for the alternate allele (Figure 3c). Four genes related to heat responses (*hsp1, hsp22, hsp101* and *ZmHsf06*) were identified, suggesting potential *cis*-regulatory elements involved in the regulation of heat responsiveness of these genes. Only limited number of *cis*-eQTL overlapped with GWAS hits and perhaps this was due to the distinct categories of SNPs could be enriched between GWAS and eQTL mapping ^37^. However, the transcription factor -- *farl8* with SNPs inside its intron regions was significantly associated with Y(II) in control specific condition and was also identified as a reGene to be regulated by a reQTL within its intron. Given the responsive difference per reGene between genotypes with reference alleles and alternative alleles, responsive patterns of reGenes could be classified into different scenarios according to the median expression value. There were 165 reGenes that exhibited stronger up-regulation response for either the reference allele (78) or the alternate allele (87) and these included examples in which both alleles were up-regulated (but to different levels) as well as examples in which only one allele was upregulated. For example, an altered InDel from ATA to TACTC in the putative 3’ UTR region of *Zm00001d050304* was associated with higher expression under heat stress (Figure 3d). Similarly, a SNP in the ~1kb promoter region of *Zm00001d046322* (Figure 3e) was strongly associated with higher heat response upregulation in genotypes with reference alleles compared to genotypes with alternative alleles. More distinct regulation patterns were observed as opposite regulations between genotypes with reference and alternative allele (Figure 3f-g).

One of our goals was to identify sequence variation that might contribute to variable gene responsiveness under heat stress. However, only the most significant eQTL SNPs were selected for performing the analysis to detect the reQTL and its associated reGene. Although the multiple testing burden could be ameliorated this way, the tested reQTL might not reflect a direct causal relationship. Haplotype analysis revealed large haplotype blocks that could be associated with variable responses of most reGenes. For example, *Zm00001d042183* was annotated as being involved in the triacylglycerol degradation pathway and we located a significant *cis*-eQTL on 1,443 bp upstream from its TSS for reQTL mapping. The analysis of all SNPs near this gene revealed multiple highly associated SNPs that exhibited high levels of linkage disequilibrium (Figure 3h). The reGene *Zm00001d030549*, that was putatively involved in the chlorophyll degradation pathway, provides another example of multiple highly associated SNPs within a large haplotype block (Figure 3i). Even though reGenes and heo-eGenes identified from our approach showed variable responsiveness between genotypes associated with different alleles, multiple SNPs exhibited similar correlations with gene expression responsiveness that fall in a large haplotype region that complicated the ability to determine the underlying causal variant.

### Changes in chromatin structure associated with heat responsiveness in maize

Chromatin accessibility provided an additional avenue to document candidate *cis*-regulatory elements under heat stress. To document changes in accessible chromatin regions with evidence of DNA binding protein footprints in response to heat stress in maize, we generated MNase-defined cistrome-Occupancy Analysis (MOA-seq) data for three biological replicates of B73 plants grown in control and heat stress (4 hrs at 40C). We initially identified peaks of accessible regions using the full MOA-seq reads. Prior work found that using only the middle 20bp of each MOA-seq read provided likely TF footprints based on known binding sites ^24^. The analysis of the high-resolution (center 20bp of each read only) MOA-seq data resulted in a set of smaller peaks (median size of 34bp compared to 179bp for full MOA-seq reads). Over 130,000 MOA-seq footprints were identified in each of the biological replicates in either control or heat condition (Supplemental Table S7 and Supplemental Data 1). TF footprints were frequently identified in regions near the TSS of many genes (Figure S8). Three biological replicates of control and heat MOA-seq samples in B73 were compared to identify 13,792 TF footprints that exhibited significant (adjusted p value <0.05) difference between conditions (Supplemental Data 1). These differential TF footprints likely revealed regions of the genome with differential occupancy of TFs or other DNA binding proteins in heat-stressed samples relative to the control samples. The analysis of genes that exhibited heat-stress inducible expression revealed examples in which there was a distinct difference in MOA-seq coverage shape (Figure 4a) as well as examples with quantitative differences in TF footprints (Figure 4b), but there were relatively few truly novel peaks that were only present in heat-stressed samples. There were 4,943 differential TF footprints located in putative promoter regions (−2,000bp from the TSS per gene) and another 1,486 located within gene regions. The differential TF footprints included more regions with increased accessibility in heat stress (11,140) compared to regions with reduced accessibility in heat stress compared to controls (2,652). The regions with increased MOA-seq coverage in heat stress compared to control might reflect increased binding or occupancy of transcription factors activated by heat stress. A metaplot of MOA-seq read depth at the heat-enriched TF footprints revealed that these regions already exhibited MOA-seq coverage in control samples but had substantially stronger signals in heat-stressed samples (Figure 4c). Heat-enriched TF footprints were detected within 2,000bp of the TSS in 8 HSFs and 18 hsps. Interestingly, 1,353 of 13,792 differential TF footprint regions were located within annotated transposable elements. Some transposable elements with heat-enriched TF footprints were also detected to be upregulated under heat stress in a previous study ^29^, such as *RLG00007Zm00001d00242 (Gypsy* retrotransposon), *RLC00002Zm00001d10560 (Copia* retrotransposon), *RLX06576Zm00001d00001* (unknown retrotransposon) (Supplemental Table S8). The regions that contained heat-enriched MOA-seq footprint peaks were used to perform motif enrichment analysis to identify potential TF binding sites. We identified 26 motifs with putative TF binding activities in the *cis*-BP database ^38^ (e.g. AP2, E2F, Myb/SANT, GATA, Dof, bHLH, NAC/NAM) and another 24 motifs that do not contain previously characterized TF binding sites (Supplemental Table S9). Prior work had shown that HSFs played important roles in regulation of gene expression in response to heat stress and variants of the HSF binding motif were often enriched near heat responsive genes ^14^. We identified an enriched sequence (GAAGCTTC) matching HSF motif -- HSFB2A -- in the heat-enriched TF footprints.

**Figure 4.**
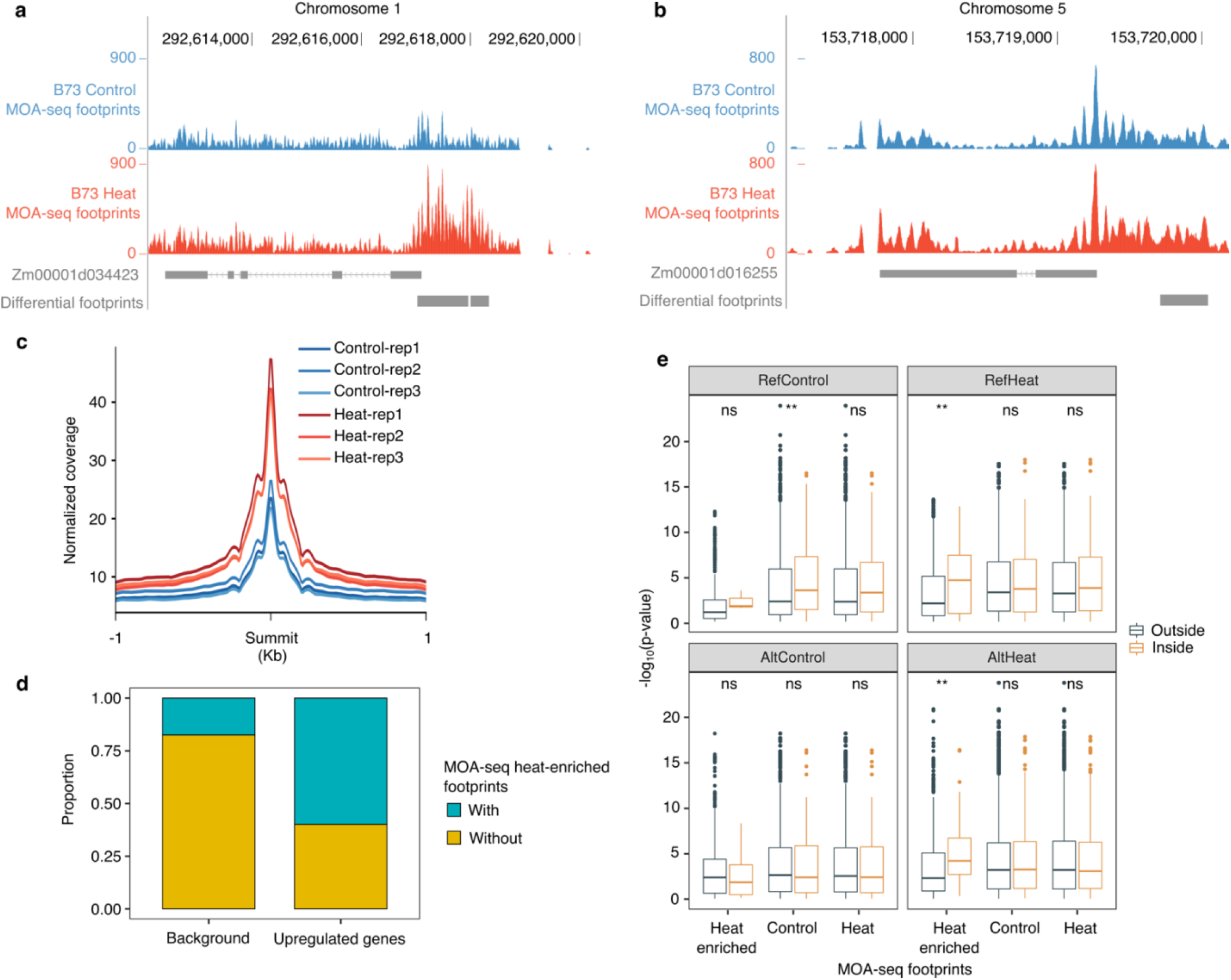
Utilization of TF footprints to identify heat response *cis*-regulatory elements. (a) An example of differential TF footprints near a B73 heat-responsive gene. Raw read coverages were normalized by RPGC (Reads Per Genomic Content) methods; (b) Same as (a), but for another heat responsive gene, *Zm00001d016255 (ZmHSF13);* (c) Normalized MOA-seq signals near 1kb extended regions of putative summits of heat-enriched TF footprints in three biological replicates under control and heat conditions; (d) Compared 793 commonly upregulated genes with 11,501 background genes for presence and absence of heat-enriched TF footprints; (e) Comparisons of interaction terms between SNPs located inside and outside of defined TF footprints located within different categories of reGenes and their 1kb flanking regions. The definition of RefControl, RefHeat, AltControl and AltHeat were the same as Figure 3c. ** indicates p-value < 1e^−2^ and ns indicates insignificant.

The same tissue samples that were used for B73 MOA-seq were also used to generate B73 RNA-seq data in the same conditions. To eliminate the experimental batch effect for comparing the changes in chromatin accessibility and potential variable regulatory elements in the panel of diverse genotypes, we focused on a subset of 793 genes that were commonly identified as upregulated genes and 11,501 genes that were commonly identified as expressed (mean CPM > 1) but not DEG in both RNA-seq datasets (the matched MOA-seq samples and the three biological replicates of B73 sampled as part of the population study). These 793 genes and 11,501 genes were classified as differentially expressed genes and control genes, respectively. We found that the differentially expressed genes exhibit significant enrichment (p-value < 2.2e^−16^) for heat-enriched TF footprints within their 2kb flanking regions compared with control genes (Figure 4d). We noted that the 2kb flanking regions of six HSFs (*ZmHsf03, ZmHsf05, ZmHsf08, ZmHsf11, ZmHsf25, ZmHsf26*) that were consistently up-regulated across genotypes contained heat-enriched TF footprints, suggesting potential regulatory regions of these HSFs. This suggested that the heat-enriched TF footprints were often associated with gene activation under heat stress. However, it was noteworthy that 42% of 793 genes that were up-regulated in heat stress did not contain heat-enriched TF footprints. These included many examples of quite strong up-regulation without evidence for altered TF footprints. The vast majority (>99%) of the 793 up-regulated genes had TF footprints within 2kb. This revealed that differential TF footprints were enriched at up-regulated genes but that some up-regulated genes had consistent TF footprints even with altered expression levels. Together, these results suggested that identified chromatin accessibility changes near heat-upregulated genes could highlight particular *cis*-regulatory elements (CREs) that might be involved in transcriptional responses to heat stress.

### Application of B73 TF footprints to identify candidate causal eQTL variants

We sought to utilize the MOA-seq data to prioritize candidate variants near the reGenes as potential causative changes. There were 258 reGenes that were expressed in many genotypes in both control and heat conditions that exhibited significant variation for heat responsiveness as well as another 518 heo-eGenes that were only expressed in heat-stressed samples that exhibited significant cis-eQTL suggesting variable activation of these genes in response to heat stress. The reGenes were classified into four groups as RefControl (n=42), RefHeat (n=78), AltControl (n=56) and AltHeat (n=87) based on which haplotype exhibited a strong response to heat stress (Alt / Ref) and whether the more responsive allele was up- (Heat) or down- (Control) regulated in response to heat stress (Figure 3c). Similarly, the heo-eGenes could be classified as RefHeat (n=313) and AltHeat (n=205) based on whether the reference or alternate allele exhibited greater expression in heat-stressed samples. The RefHeat or AltHeat groups of reGenes or heo-eGenes all exhibited up-regulation in response to heat and many of these genes (30-37%) have a heat-enriched MOA-seq footprint within 2kb of the gene (Supplemental Table S10). These were enriched relative to other expressed genes. In contrast, the RefControl and AltControl genes that did not necessarily show up-regulation in response to heat stress have only 14-18% of genes with a heat-enriched MOA-seq footprint, which was similar to all expressed genes.

In most cases, there were multiple variants that were significantly associated with the heat response (for reGenes) (Figure 3h-i) or expression levels in heat stress samples (for heo-eGenes). We assessed whether at least one of the most highly associated (top 3 p-values) variants was located within a heat-enriched MOA-seq footprint for the subset of genes that contained these footprints. Among the genes that had decreased expression in response to stress (RefControl or AltControl) with a heat-enriched MOA-seq footprint there were only 12.5% with a highly associated variant located within the small footprint region. In contrast, 24% of the RefHeat and 30% of the AltHeat reGenes with heat-enriched TF footprints had at least one highly associated variant that was located in the footprint region (Supplemental Table S10). There were 21.7% and 22.6% of the RefHeat and AltHeat heo-eGenes, respectively, that had a highly associated variant within the heat-enriched TF footprints. Through rerunning the reQTL mapping using all variants located in reGenes and their flanking regions, it revealed that significant difference of interaction effects for variants located inside of heat-enriched TF footprints and variants outside of the TF footprints for either RefHeat (Wilcox-test, p=8.67e^−3^) or AltHeat reGenes (Wilcox-test, p=2.07e^−3^; Figure 4e; Supplemental Table S11). For both reGenes and heo-eGenes, our findings suggested a robust power of heat-enriched MOA-seq signals derived from a single genotype on predicting responsive genomic regions. It was worth noting that in many of the AltHeat genes there was a significant response to heat stress for both the reference and alternate allele, but the alternate allele had a significantly stronger response. In these cases, the B73 allele might still have a heat-enriched footprint and the alternate allele might have a variant that allows for enhanced TF binding. Together, our results suggested overlaps between heat-enriched TF footprints and identified responsive variants might pinpoint regulatory genomic regions in maize under heat stress.

### Transient expression assays confirm allelic variation for heat response

We were interested in determining whether the 2kb proximal regions of the reference and alternate alleles of reGenes would be sufficient to recapitulate the variable responses to heat stress. We selected three pairs of alleles -- *Zm00001d005114* (B73) / *Zm00014a020759* (Mo17), *Zm00001d017187* (B73) / *Zm00035ab255850* (MS71) and *Zm00001d042183* (B73) / *Zm00039ab143500* (Oh43) for these experiments. Two of these examples exhibited significant differences for TF footprints in response to heat stress (Figure 5a-b) while the other gene had ~19.4% increased signals in the TF footprints under heat stress but did not pass the significance filter (Figure 5c). The two genes with significant differences in the footprints *Zm00001d005114* and *Zm00001d017187* showed differential response to heat stress for the reference and alternate alleles (Figure 5d-e). For both of these genes the reference allele had a much weaker response to heat stress as compared to the alternate allele. The third gene, *Zm00001d042183*, exhibited no significant response to heat stress in genotypes carrying the reference allele, but had significant up-regulation in genotypes with the alternate allele (Figure 5f). A comparison of the allele-specific expression in three biological replicates of B73xMo17 or B73xOh43 F1 hybrids was used to validate the allelic ratio in control and heat stress plants for *Zm00001d005114* and *Zm00001d042183* that were heterozygous for the reference and alternate alleles in these hybrids (Figure 5g-h). Both of these genes showed a significant difference in the ratio of reference:alternate allele expression in heat-stress samples compared to plants grown in control conditions, confirming the *cis*-allelic variation for heat responses for these alleles.

**Figure 5.**
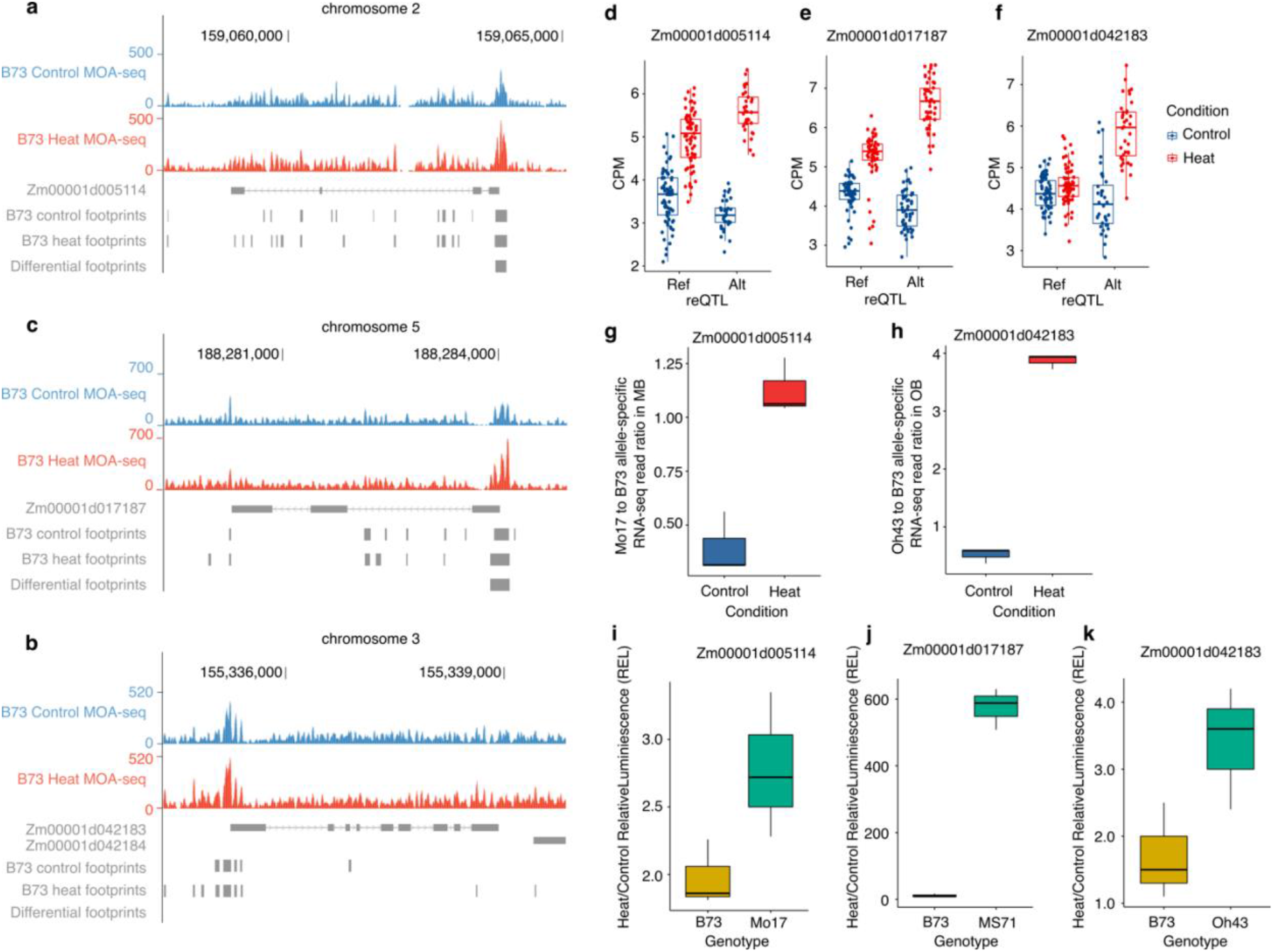
Validations of allele-specific regulatory genomic regions for reGenes. (a-c). The genome browsers showed MOA-seq signals of reGenes between heat and control in B73 genome coordinates. MOA-seq genome coverage was normalized using the RPGC methods to make tracks comparable. (a) *Zm00001d005114;* (b) *Zm00001d017187;* (c) *Zm00001d042183;* (d-f) Expressions of assessed reGenes in genotypes associated with the detected reQTL as reference allele or alternative allele under control and heat condition. (d) *Zm00001d005114;* (e) *Zm00001d0171873;* (f) *Zm00001d04218;* (g-h) Raw read ratios of (g) Mo17 or (h) Oh43 specific allele to B73 specific allele of *Zm00001d005114* or *Zm00001d042183* in the hybrid MB or OB under control and heat condition (n=3); (i-k) Around 2kb putative promoter region per reGene was amplified separately from B73 genotypes and relative alternative genotypes for the DUAL-luciferase assay (n=3). (i) Promoters amplified from *Zm00001d005114* and *Zm00014a020759;* (j) Promoters amplified from *Zm00001d017187* and *Zm00035ab255850;* (k) Promoters amplified from *Zm00001d042183* and *Zm00039ab143500*.

The 2kb promoter sequences (including the transcription start site and 5’ UTR) were cloned and fused to a luciferase reporter construct for assessing segregations of their activation roles for reGenes under heat stress. Alignments of the two promoters revealed significant conservation in the region immediately upstream of the transcription start site, but in each case the more upstream regions were not alignable, often due to polymorphic transposon insertions (Figure S9). Two of the examples included a differential heat-enriched MOA-seq footprint that overlaps the transcription start site (Figure S9a-b). The third example had several MOA-seq footprints that were only detected in heat-stressed samples but these were not identified as significantly heat-enriched (Figure S9c). Each of the three pairs of promoters exhibited differences in the level of heat activation using the DUAL luciferase reporter assay that mirrored the RNA-seq data for the reference and alternate alleles (Figure 5i-k). In each case the reference allele had a much smaller response to heat relative to the alternate allele. It is worth noting that the level of activation in response to heat stress was highly variable for the three different promoters with *Zm00001d017187* exhibiting much stronger heat response than the other two promoters.

## Discussion

Plants frequently encounter abiotic stresses and there is evidence for variable stress responses in many species ^39^. This study focused on characterizing the sources of natural variation for gene expression responses to a heat stress event. Our characterization focused on a moderate (non-lethal) 4 hour heat stress treatment at 40C. While many genes respond rapidly (<30 minutes) to heat stress there are quite dynamic changes in the response at early time points ^14^. In order to minimize potential complications due to sampling over a ~15 minute period for a full population, we focused on an intermediate 4 hour heat stress treatment. There are many genes with highly conserved responses to a heat stress in the population that we surveyed. Many of the genes with strong, consistent responses to the heat stress event are heat stress proteins that likely act as chaperones. This confirms that the panel of genotypes that was used in this study were all able to generate a strong heat response.

Gene expression levels were highly variable within the population that was used for this study. Prior studies have documented many examples of *cis*- and *trans*-eQTL in maize ^40–43^. We find similar trends in terms of the proportion of *cis*- and *trans*-eQTL as well as the relative magnitude of effects. Relatively few major *trans*-eQTL hotspots were identified in our analysis. There are a handful of genomic loci that influence expression of 10-20 other genes but no loci with hundreds of *trans*-eQTLs. Given that our population consists of maize inbreds that have been subject to artificial selection and are phenologically restricted this may not be all that surprising. It is also likely that our experiment was under-powered to detect *trans*-eQTL ^44^.

### Mapping sources of cis-variation for heat stress response

A primary goal of our study was to document variation in *cis*-regulatory elements that provide variable gene expression responses to heat stress. In order to document genes with *cis*-eQTL for heat responses, we implemented response eQTL analysis that were largely adapted in human studies ^45–47^. Other than assessing gene responsive variations between a few genotypes, our approach employed hundreds of genotypes to substantially increase the power of statistical testing for pinpointing potential responsive regulatory regions and avoid transcriptomic instability of a single genotype in response to external heat stress stimuli. This approach identified 258 reGenes that exhibit differential responses to heat stress. While this is a promising approach to document genes with significant variation in responsiveness that is associated with nearby genetic variation, there are a class of genes that can not be identified using this approach. The set of genes that do not exhibit detectable expression in control conditions that are activated by heat stress (heat stress expressed only by genes) can not be assessed for variation. We found another set of 518 heat-expressed only genes that have *cis*-eQTLs in heat stress, suggesting differential activation in response to the heat stress. These two sets of genes represented examples of *cis*-variable responses to heat stress. We found similar numbers of genes for which the reference or alternative allele exhibited a strong response to the heat stress and the genes include examples of both variable activation and variable repression in response to heat stress. It is worth noting that similar to all GWAS approaches there are limitations in detection of rare alleles that exhibit variable responses. We restricted our analyses to sequence variants with a minor allele frequency of at least 0.10 in this panel. However, the reQTLs that exhibited significant effect had a median minor allele frequency as 0.33.

### Applying TF footprinting and transient assays to characterize cis-variable responses

The approaches to document genes with variable response often focus on using the most highly associated sequence variant in either control or heat expression data. However, many studies have shown that there are often multiple sequence variants present together in a haplotype with high levels of linkage disequilibrium ^48^ and it can be quite challenging to resolve which of these variants are causal for the expression difference. The majority of reGenes and heo-eGenes have multiple nearby genetic variants that are highly associated with variable expression responses. In order to develop prioritized candidate variants that might be responsible for the variable stress responsive expression, we utilized a recently developed method 24 to perform genome-wide transcription factor footprinting searching.

MOA-seq was performed for B73 in both control and heat stress conditions. This approach identifies ~100,000 regions with TF binding for each replicate in either of conditions. However, ~60% peaks commonly presented in both control and heat TF footprints using the merged data. A comparison of the control and heat stressed samples using replicates identified over 13,000 TF footprints with significant variation in heat stress and control conditions with ~80% of these exhibiting increase in heat compared to control. Genes that are up-regulated in response to heat exhibit a significant enrichment for TF footprints that are significantly increased in heat stress. However, there are several observations that should limit the simplistic view of novel TF binding being associated with heat responsive gene expression. First, we found very few examples of truly novel TF footprints in heat stress compared to control. Instead the majority of significant differences in MOA-seq data represent examples in which there is quantitative variation such that a footprint is present in control conditions but becomes stronger in heat stress tissues. This likely suggests changes in average TF occupancy in a cell population following heat stress rather than novel binding events that are not detectable in control conditions. Similar observations were made regarding chromatin accessibility changes in rice plants subjected to varying water availability ^49^ or in maize plants subjected to heat stress conditions ^50^. Second, there are many genes that are up-regulated in response to heat stress, including some that are only expressed in response to heat, that do not have evidence for variable TF footprints in surrounding regions. Many of these genes have consistent TF footprints in control and heat stress samples. This may suggest that TF occupancy does not change in response to heat stress. Instead, there may be variable activity of the TFs due to post-translational modifications that provide altered regulation.

The MOA-seq data generated from B73 provided insights into one allele that was present in our population. When B73 is the more responsive allele we can use the MOA-seq data to prioritize potential genetic variants that exhibit significant association with expression responses. All of the reGenes for which B73 has the more responsive allele have at least one highly associated SNP or InDel within their 2kb flanking regions that is located within a TF footprint. A subset of these are located within TF footprints that exhibit significant differences between heat stress and control samples.

In order to provide further evidence for causal variants that influence heat responsive gene expression it is necessary to perform functional assays. We developed a transient protoplast expression system to assess the heat responsive activity of promoters. We initially focused on utilizing the 2kb promoter region for the reference and alternate haplotypes to drive expression of a reporter. The dual luciferase assay was then used to compare the relative expression in control and heat stressed protoplasts. We found that we could recapitulate the allelic variation observed in plants in these assays. In the examples tested to date we have used the full haplotype of the promoter region and this has included multiple sequence variants. However, this assay can be further utilized to perform targeted sequence changes to monitor the functional impact for each of the specific sequence variants. This system will be useful as we seek to document the molecular basis allelic variation in gene expression responses to a heat stress event.

Heat stress is one of the abiotic stresses that plants encounter regularly. In this study we have developed approaches to document and characterize natural genomic variation that can regulate heat stress responses of genes at the transcriptomic level. Changes in gene expression are one important mechanism plants utilize to survive abiotic stresses whereas we have limited understanding of the molecular mechanisms that generate natural variation for allelic responses. Our study provides further insights into the prevalence and characterizing sources of natural variation for gene expression responses to heat stress in maize or other plant species.

## Materials and Methods

### Plant materials, growth and treatment

The SNP set of 509 inbreds from the WiDiv (Wisconsin Diversity) Panel was retrieved from a previous publication ^31^ for genotype subtraction. Biallelic SNPs were retained upon multiple thresholdings (no missing data across 509 inbreds; MAF > 0.05; heterozygosity <= 0.2). To reduce the computational cost, we randomly selected 1,000 SNPs per chromosome for each inbred. The entire set of selected SNPs were used to calculate Euclidean distances between inbreds. The Ward’s method of hierarchical clustering was then employed to determine the initial 120 clusters. One representative genotype was subtracted from each of 120 clusters. All genotypes were grown in the growth chamber for 14 days in 30 C/20 C 12h/12h day/night cycle. Positions of plants in the growth chamber were randomly shuffled during the plant growth to minimize the microenvironmental effect. On the 14th day, plants subjected to heat stress were treated for 4 hrs under 40 C and plants from corresponding genotypes were under control conditions of 30 C in parallel. Control and heat environments were separately implemented in two identical growth chambers. Once the heat treatment was completed, the third leaf per plant was collected and 2-3 plants per genotype were merged to represent one genotype for RNA-seq data generation in each condition. Three biological replicates of B73 were separately inserted into the panel during sample collections for checking experimental repeatability.

### Phenotypic measurements

The same set of genotypes used in RNA-seq data collection with three replicates were subjected to the same control and heat conditions for chlorophyll fluorescence measurement. Chlorophyll fluorescence parameters including photochemical and non-photochemical quenching were analyzed using a pulse-amplitude modulated chlorophyll fluorescence camera (MAXI Version IMAGING-PAM M-Series, Heinz Walz, Germany) with a blue (450 nm peak λ) actinic and measuring light-emitting diode (LED) array and an IMAGE-K7 CCD camera with a Cosmicar-Pentax zoom objective lens. All data was acquired in a dark room. Whole-plant samples were dark adapted for 15 min, the third leaf was detached for a given sample and was directly taped to a non-fluorescent background underneath the LED array with near-infrared LEDs on only. Zoom and focus were adjusted to acquire data on four samples in one image, a control and heat treated sample from two different genotypes. The camera aperture was set to 1.0 for all samples and the measuring light intensity and gain were at settings of 2 and 3 respectively (frequency = 1.0) to maintain a current fluorescence yield (F_t_) between 0.100 - 0.180. Once F_t_ came to equilibrium (10-20 seconds after turning on the non-actinic blue light source) a saturation light pulse was applied and the minimum fluorescence yield (F_o_) and maximal fluorescence yield (F_m_) were recorded. Samples were then exposed to 3 min of actinic light of approximately 950 μmol m^−2^s^−1^ to determine photochemical Y_II_ and non-photochemical quenching parameters Y_NPQ_. The other non-photochemical loss Y_NO_ was determined using the equation of Y_II_ + Y_NPQ_ + Y_NO_ = 1 ^51^. Data was collected within an area of interest outlining the entire leaf. The maximum value of Y_NPQ_ at two minutes following actinic light treatment was used for treatment comparisons. Due to the capacity of measurement, the entire genotypes were split into 4 blocks using a split block experimental design with B73 as checks in each block under either control or heat condition. BLUPs (Best Linear Unbiased Predictions) of Y_NPQ_, Y_II_ and Y_NO_ were separately estimated for control and heat data using a linear mixed model with all effects (genotype, block, replicate) treated as random. In addition, the broad-sense heritability (H^2^) of each chlorophyll fluorescence parameter per condition was estimated as the proportion of genotypic variation to the total variation.

### Genome-wide association study

The same set of filtered genomic variants in the studies and BLUPs estimated from phenotype data were used for genome-wide association study (GWAS). The GWAS association model FarmCPU was employed using rMVP v1.0.6 ^52^ to detect genotype-phenotype association signals. The effective number of variants was determined as 453,641 using GECv2.0 ^53^ with default settings. Two levels of bonferroni correction were applied to set the cut-off at 1.10e^−7^ (0.05/453,641) and 2.20e^−6^ (1/453,641).

### RNA-seq data generation and data processing

RNA-seq data were generated using NovaSeq 6000 in paired-end 150bp mode. Raw reads per library were preprocessed using Trim-galore (Babraham Bioinformatics) with default settings. Preprocessed reads were aligned against the indexed B73 AGPv4 reference genome ^54^ using HISAT2 ^55^. Alignment files were sorted and indexed using SAMtools (v1.9) ^56^. The longest transcript was used to represent the individual gene model in the B73 AGPv4.41 version (downloaded from Ensemble). Raw counts per gene were calculated using HTSeq-count (v0.11.2) ^57^ and normalized into CPM (Counts Per Million mapped reads) value per gene by library sequencing depth using DESeq2 ^58^. Genes with absolute log_2_ fold change greater than 1 and adjusted p-value less than 0.05 were considered as differentially expressed genes (DEGs) in B73 samples.

To confirm identities of accessions employed in this study, we employed the Broad’s Genome Analysis Toolkit (GATK) -- “RNA-seq short variant discovery” pipeline to call SNPs using alignment files generated from HISAT2. Comparing nucleotides on same SNPs generated from RNA-seq and whole-genome resequencing data of identical accessions, we retained accessions with a consistent SNP rate greater than 80% in the study, resulting in 102 genotypes for further analysis.

### Comparing gene expressions between control and heat conditions across genotypes

Genes with CPM > 1 in at least 10% genotypes in the panel in either control or heat condition were retained for the analysis. A paired wilcoxon test was performed to compare expressions between control and heat conditions. The p-value per gene was adjusted for multiple testing corrections using the Benjamin-Hochberg methods. Genes with adjusted p-value < 0.01 were considered statistically significant. Significant genes with log_2_ transformed heat/control greater than 1 were considered as up-regulated genes and less than 1 were considered as down-regulated genes. Genes with the median value of log_2_ transformed ratio of heat to control greater than 0 and 90% quantile of the log_2_ transformed ratio of heat to control greater than 0 were considered as genes with consistent upregulation in the population.

### GO enrichment analysis

Upregulated genes identified in B73 replicates inserted into the panel transcriptomic experiment were considered as the tested gene set. GO terms per gene were retrieved from the maize GAMER dataset ^59^ for maize AGPv4 gene models. GOATOOLS ^60^ was implemented to perform the GO enrichment analysis. Any GO term with a bonferroni corrected p value less than 0.05 was considered significant.

### Genotyping and eQTL mapping analysis

Missing genotype SNP calls of 509 inbreds were imputed using Beagle (version 05.Apr.21) ^61^. SNPs of the subset inbreds consisting of 102 genotypes employed in this study were subtracted and retained with Minor Allele Frequency (MAF) >= 0.1 and heterozygous rate <= 0.1. Highly correlated SNPs were pruned using PLINK ^62^ (--indep-pairwise 50 5 0.99). InDels from the same prior study were retrieved and only InDels with no missing data on studied genotypes were retained to merge with filtered SNPs together as the final set of 1,132,322 genotype variants. The first five principal components (PCs) were calculated using the R function prcomp. Each gene with CPM > 1 in at least 10% genotypes was retained for analysis in either control or heat condition. CPM values per gene under control or heat condition were transformed into a normal distribution using the R package bestNormalize (https://github.com/petersonR/bestNormalize) separately. Twenty-five hidden factors were separately inferred from control and heat expression data using PEER ^63^ for controlling variance. SNPs located nearby 1Mb of gene regions were classified as *cis*-eQTLs, otherwise as *trans*-eQTLs. The R package MatrixeQTL ^64^ was used for eQTL mapping in controlling 5 PCs and 25 hidden factors in either control or heat conditions. SNPs with Benjamini-Hochberg corrected p-value < 0.01 in each condition were considered as eQTLs. Genes with significantly associated eQTLs were considered as eGenes. The proportion of variance explained by each identified eQTL was calculated based upon the MatrixeQTL manual.

### Trans-eQTL hotspots identification

*Trans*-eQTLs identified by eQTL mapping in each condition were used for detecting *trans*-eQTL hotspots. The maize genome was equally segmented into 10kb bins. Any SNP within each 10kb bin targeted greater or equal to 3 genes remotely (>1Mb) was considered as candidate *trans*-eQTL. Each 10kb bin with more than 10 targeted unique gene sets was considered as a *trans*-eQTL hotspot. Based on genes targeted by *trans*-eQTLs in control and heat conditions, we split the number of targeted genes by *trans*-eQTL hotspots into common targeted genes by two conditions, uniquely targeted genes by control and uniquely targeted genes by heat.

### Response eQTL (reQTL) detection

We modified a prior published method ^47^ for priorizing candidate *cis*-eQTLs for reQTL detections in this data. Specifically, we focused on genes that were commonly expressed in both of conditions and narrowed candidate *cis*-eQTLs for reducing multiple testing burden based on following criteria: 1) For each eGene per condition, we selected the top significant *cis*-eQTL. If multiple top significant *cis*-eQTLs existed, we randomly picked one for testing; 2) For eGene with the identical *cis*-eQTL in both conditions, we retained it for testing; 3) For eGene with non-overlapped *cis*-eQTLs between two conditions, we picked one *cis*-eQTL randomly if two *cis*-eQTLs from control and heat are in high LD (> 0.8), or we retained both of *cis*-eQTLs if they are in low LD (<= 0.8); 4) For eGene with only detected *cis*-eQTL in one condition but not the other, we retained this *cis*-eQTL for the analyis. Selected candidate *cis*-eQTLs were input into a linear mixed model below for detecting reQTLs using the R package lmerTest ^65^:

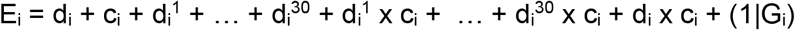

where i was the i^th^ sample, E_i_ was considered as the normalized expression value for the i^th^ sample, d_i_ was the *cis*-eQTL allele dosage for i^th^ sample, c_i_ was experimental condition for the ith sample (control or heat), d_i_^1^ to d_i_^30^ represented 30 covariates employed in this study (5 principal components + 25 PEER hidden factors), d_i_ x c_i_ was the interaction between *cis*-eQTL and condition and (1|G_i_) was the random effect of individual genotype. The p-value of the interaction term was estimated using the Satterthwaite and Kenward-Roger methods and then adjusted using the Bonferroni methods. The *cis*-eQTL relative to its targeted gene with adjusted p-value less than 0.01 were considered as reQTLs.

### MOA-seq library construction and data processing

To assess the changeable chromatin accessibility under heat stress, we collected leaf tissues from plants grown in the same growth stage under the same control and 4 hour 40C heat conditions as the large panel experiment. Collected tissues were processed for MOA-seq libraries constructions following a previous protocol ^24^. For B73 samples, the genome index of the reference genome was built using STAR (v2.7.9a) ^66^. Raw reads of MOA-seq data were preprocessed using SeqPurge (v2019_09) ^67^ with parameters of “-min_len 20 -threads 20 -qcut 0 -prefetch 50000”. Overlapping paired reads were merged into single reads using FLASH (v1.2.11) ^68^. Processed reads were aligned against the indexed reference genome using STAR (v2.7.9a) ^66^. As STAR is designed to map RNA, we set the flag -- alignIntronMax 1 for DNA, this presence and Introns from being allowed / expected by STAR. Alignment files were converted into bam formats using SAMtools (v1.9) ^69^. Alignment fragments with less than 81 bp and MAPQ as 255 were retained for further analysis. To generate high resolution maps, each read was shortened to 20bp centered around the middle of the read. Read shortening was performed using awk: for reads with uneven number of bases, the middle base was taken and then read extended 10bp to each site. For reads with even numbers of bases, one of the two middle bases was chosen randomly and the reads were extended 10bp to each site. Each filtered alignment file was converted into bigwig format using bedtools v2.29.2 ^70^. The effective genome size was determined using the k-mer size of ^60^. Coverage of alignment files were normalized to visualize using bamCoverage RPGC (Reads Per Genome Coverage) normalization function. MOA-seq TF footprints were determined using the “macs3 callpeak” function (https://github.com/macs3-project/MACS) with parameters of “-s 20 --min-length 20 --max-gap 40 --nomodel --extsize 20 --keep-dup all --buffer-size 10000000”. Distance of TF footprints to TSS was calculated using annotatePeak.pl function ^71^. We then employed the “DiffBind” (v2.12.0) function in R ^72^ to detect differential footprints between control and heat in B73 samples with FDR < 0.05.

### Motif discoveries

Summits of heat-enriched TF footprints were redefined using the “macs3 refinepeak” function and merged B73 replicates in heat condition. For each summit, flanking regions were extended according to the median length of TF footprints and nucleotide sequences were extracted from the B73 AGPv4 genome in the given region. Identified heat-enriched MOA-seq footprint sequences were input for STREME (v5.4.1) ^73^ to identify enriched motifs with shuffled genomic sequences as the background. Identified motifs were compared with PWMs (Position Weight Matrix) in cis-BP public TF database ^38^ using tomtom (version 5.0.5) ^74^. Searched motifs with p-value less than 0.05 were considered as significant and only the most significant motif matching the database was considered as the target motif for the query PWM.

### RNA-seq data generation and processing in maize hybrids

To evaluate allele-specific gene expression, we generated RNA-seq data of two selected maize hybrids including Mo17 x B73 and Oh43 x B73 grown together with B73 samples for MOA-seq data generation. All parental lines were also included in our heat stress panel experiment. SNPs between two alleles in one hybrid were retrieved from the raw SNP sets employed in the subpanel. The SNP set per hybrid was separately employed to mask the B73 AGPv4 reference genome using bedtools v2.29.2 ^70^. Raw RNA-seq reads generated from each hybrid were aligned to the respective masked reference genome in B73 AGPv4 coordinates using the same steps as above. Allele-specific alignments per sample were then split using the SNPsplit (v0.5.0) ^75^ and allele-specific read counts were calculated for each split alignment file using HTSeq-count (v0.11.2) ^57^.

### DUAL luciferase reporter assay for candidate reQTLs/reGenes

We amplified 2kb regions upstream of the start codon for reGenes pairs that displayed differential heat responsiveness between alleles. These regions were cloned into a minimal vector driving a Firefly Luciferase reporter through a combination of NEBuilder HiFi Assembly (E5520S) and traditional restriction enzyme/ligation-based approaches. Both approaches leveraged a backbone double-digested with SpeI-HF (R3133S) and SacI-HF (R3156S), dephosphorylated (Quick CIP, M0525S), and gel extracted (Omega E.Z.N.A. Gel Extraction kit, D2500-01) prior to ligation or HiFi assembly. HiFi reactions were designed and conducted as suggested in the product manual. Traditional clones were generated by amplifying regions of interest and creating corresponding SpeI/SacI restriction enzyme sites. PCR products were then purified (Zymo DNA Clean & Concentrator-5 kit, D4013), double-digested with SpeI/SacI, and ligated (T4 DNA Ligase, M0202S) into the prepared backbone. Colonies were screened and sequenced through Primordium plasmid sequencing (www.primordiumlabs.com) prior to further use.

The conditional reporter was co-transformed into maize leaf protoplasts alongside a Renilla Luciferase driven by a constitutive 35S reporter in three separate transformation events, using ~200k protoplasts per transformation. Protoplasts were generated as previously described ^76^, were transformed with a total of 10ug of plasmid DNA (8ug conditional reporter, 2ug constitutive reporter), allowed to recover for 16 hours following transformation, and then evenly split and either subjected to a +10C heat stress in a water bath or left at room temperature for three hours. Protoplasts were spun down and snap frozen at the end of the stress event. Dual Luciferase Assays were conducted per manufacturer instructions (Promega Dual-Luciferase Reporter Assay System, E1960), and measurements were taken on a Promega GloMax Explorer Plate Reader (GM3500).

## Supplemental Information

### Supplemental Figures

**Figure S1.**
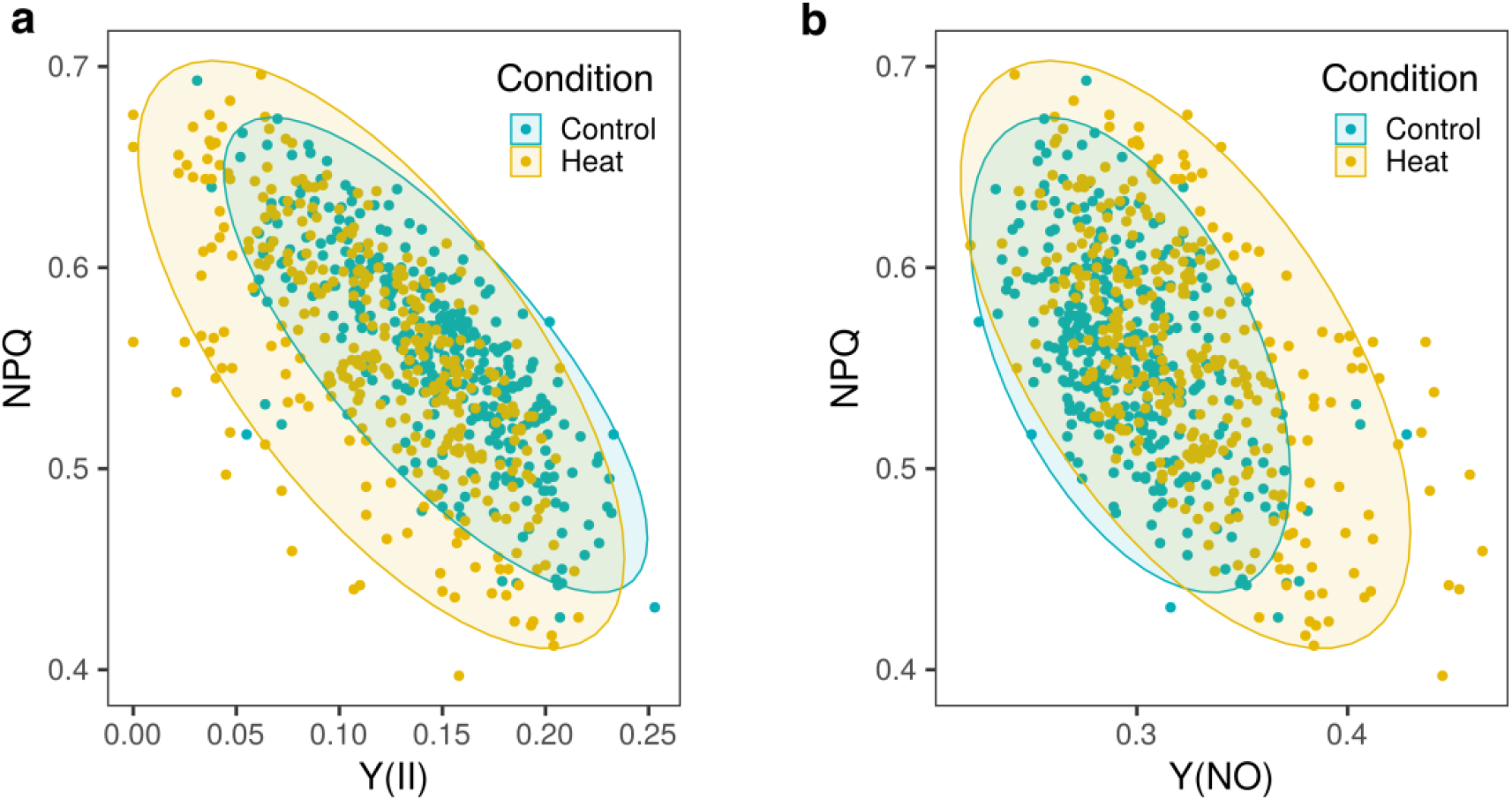
Segregation between heat stressed plants and control plants using measured photosynthetic parameters. (a) Heat stressed plants and control plants were segregated using Y(NPQ) and Y(II); (b) Heat stressed plants and control plants were segregated using Y(NPQ) and Y(NO).

**Figure S2.**
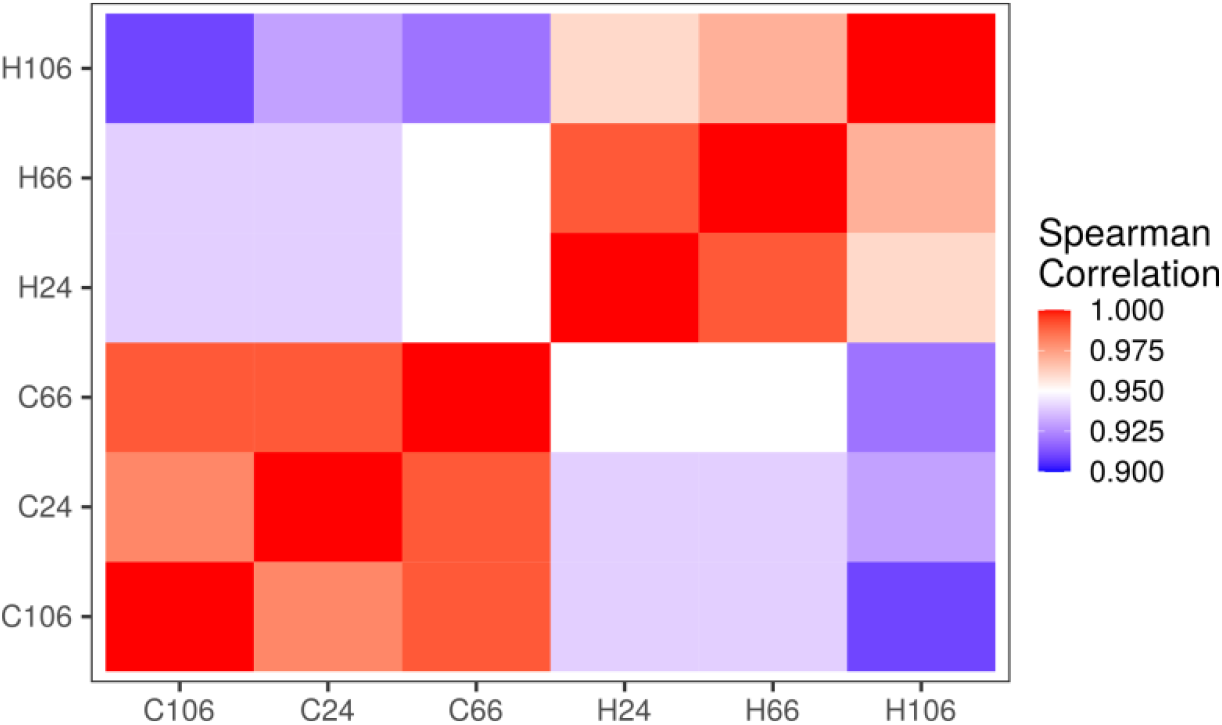
Pairwise correlations between B73 samples employed in the large panel heat stress using normalized gene read counts. Spearman correlation was employed for testing. Genotype code 24, 66 and 106 represented three biological replicates of B73 in both control (C) and heat (H) conditions.

**Figure S3.**
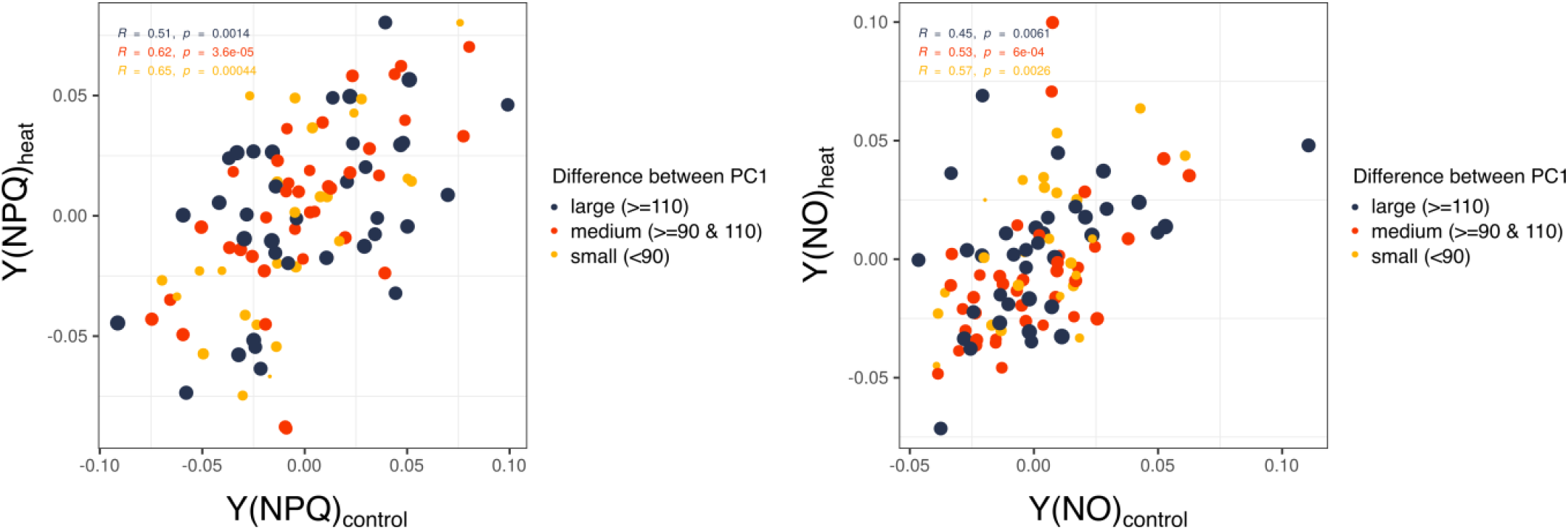
Correlations between phenotypic and transcriptomic data. Genotypes with variable levels of PC1 differences were split into three classes: large, medium and small. The correlation of BLUPs generated from Y(NPQ) and Y(NO) in control and heat condition was separately calculated.

**Figure S4.**
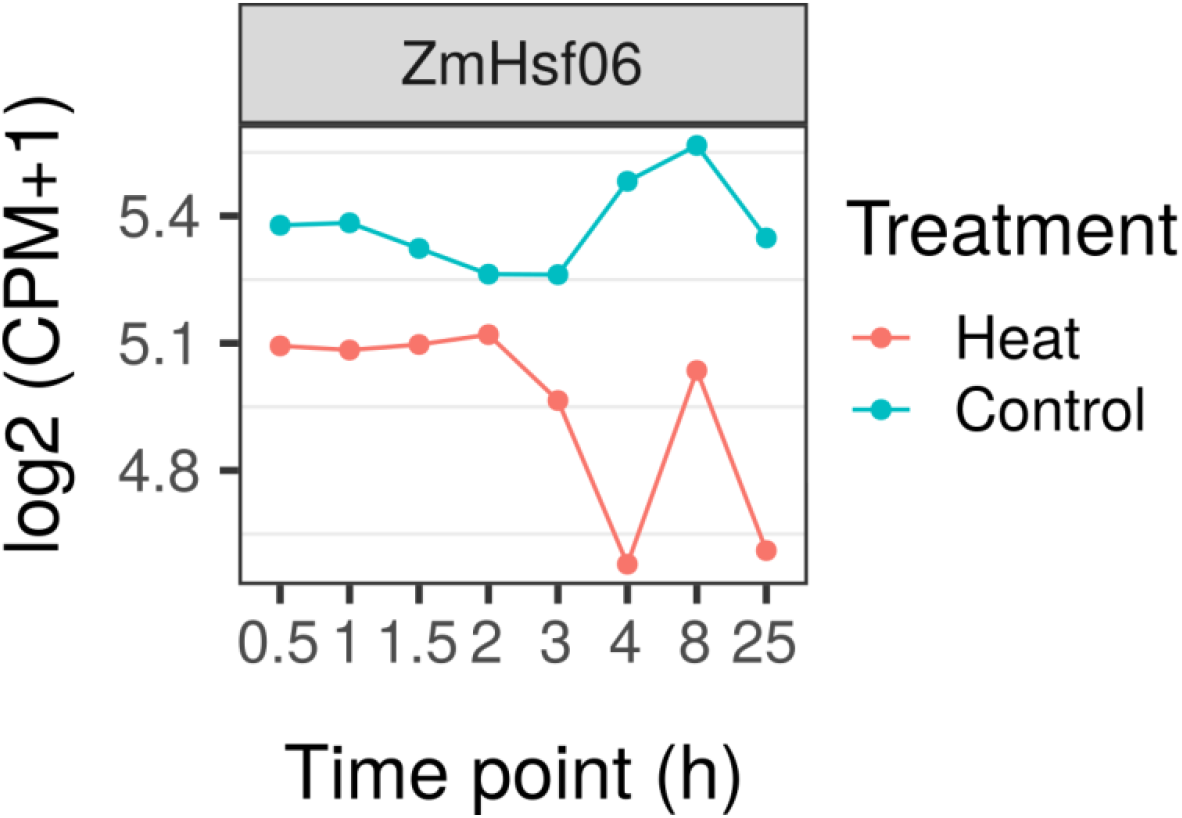
Time-series expression of ZmHsf06 under control and heat conditions in six different time points. Raw RNA-seq read counts were retrieved from a previous published paper ^14^. Expression value was normalized to counts per million mapped reads (CPM).

**Figure S5.**
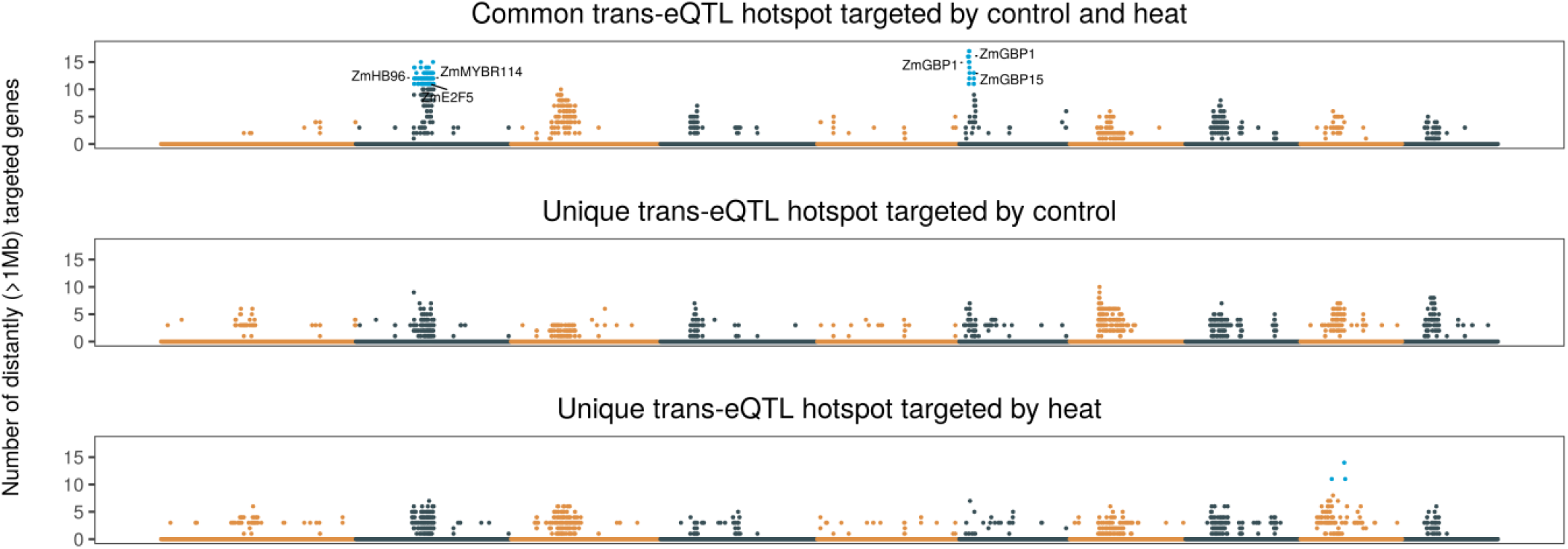
*Trans*-eQTL hotspots identified in this study. The genome was segmented into a 10kb bin. Each dot represents one 10kb and any SNP within each bin containing >= 3 targeted genes was retained. Any bin with more than 10 distantly targeted genes was considered as a *trans*-eQTL hotspot (labeled as blue). Known transcription factors were labeled for *trans*-eQTL hotspots.

**Figure S6.**
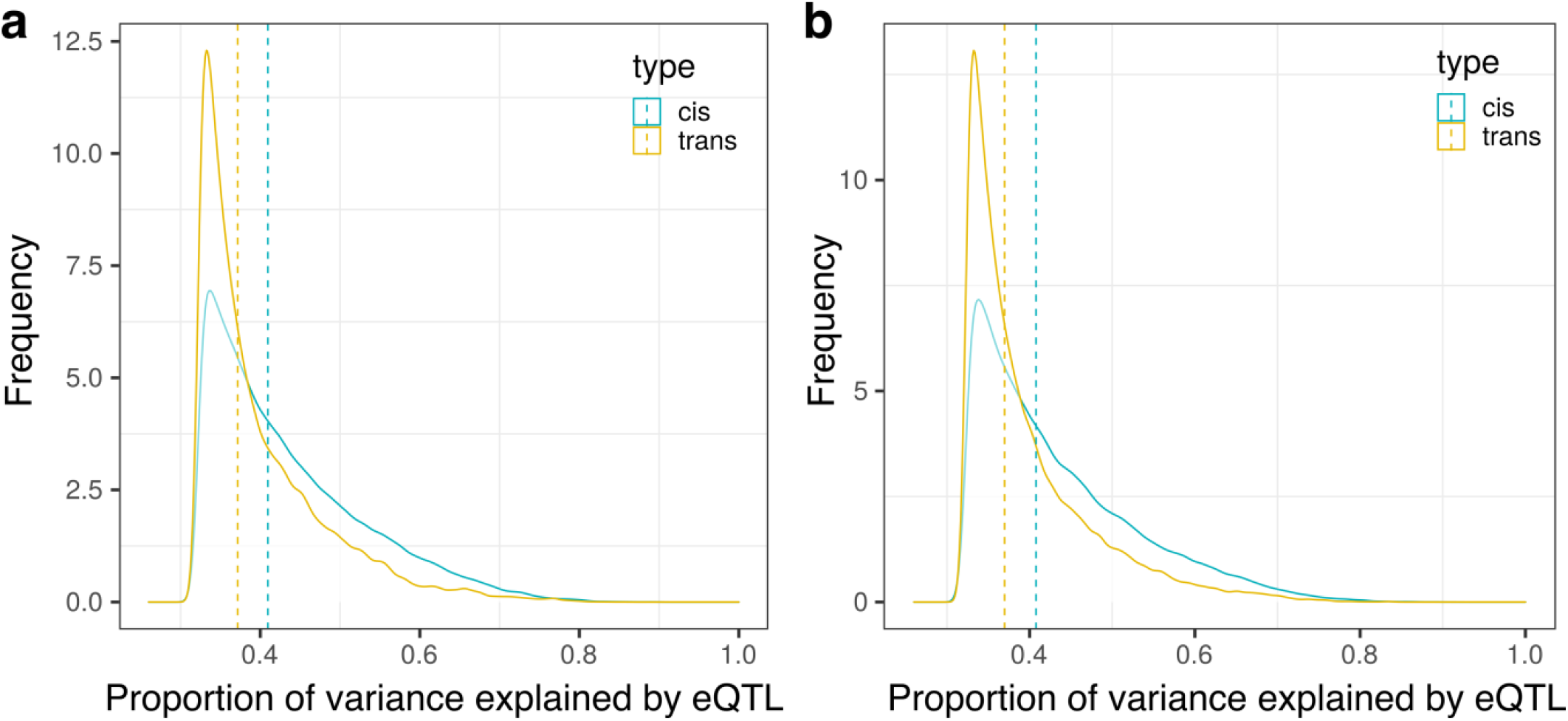
Distributions of proportions of variance explained by *cis*-eQTLs and *trans*-eQTLs identified using (a) control and (b) heat expression data. Dashed vertical lines indicate median value of either *cis*- or *trans*-eQTLs.

**Figure S7.**
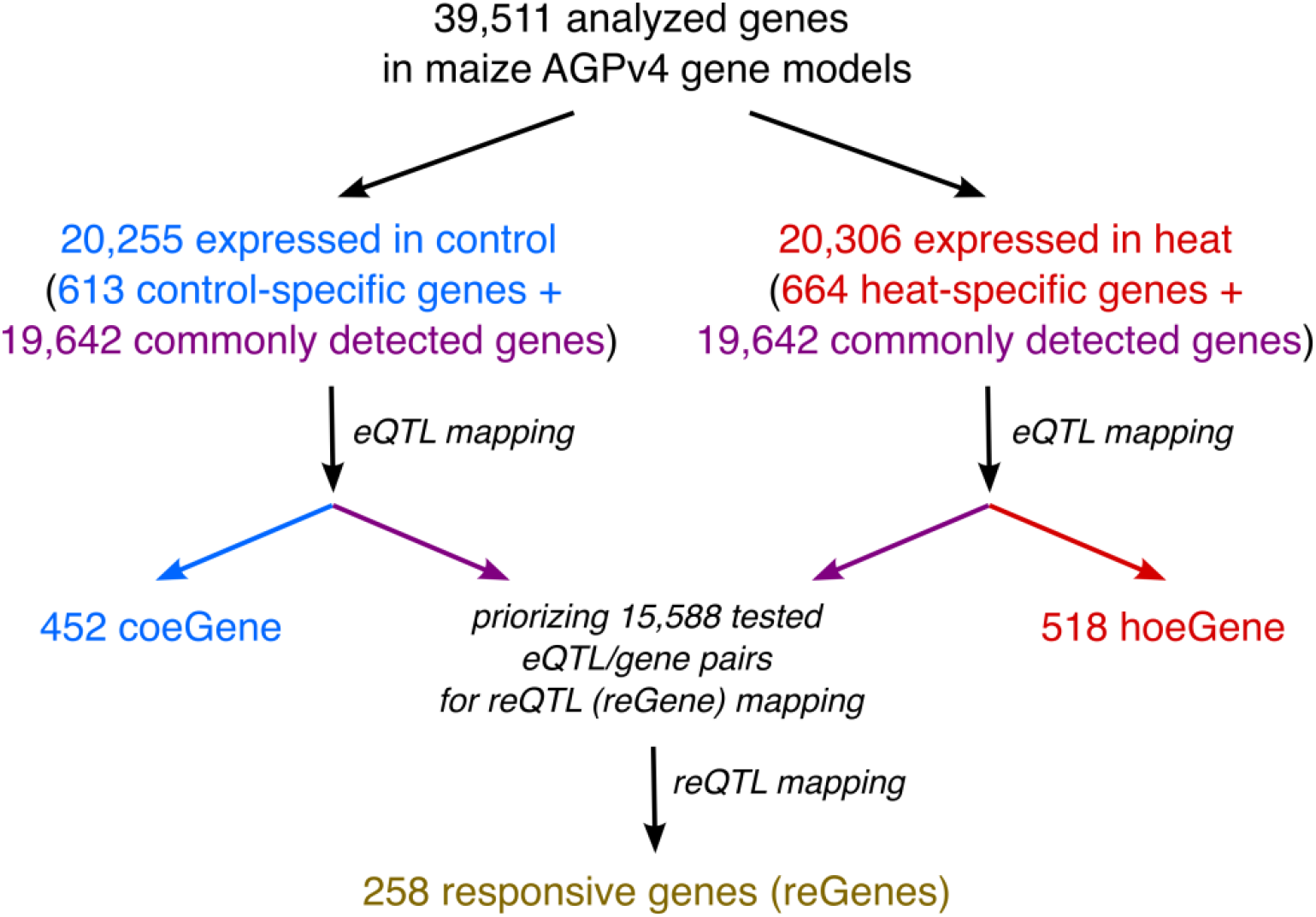
Workflow for identifying reGenes and eGenes in control and heat.

**Figure S8.**
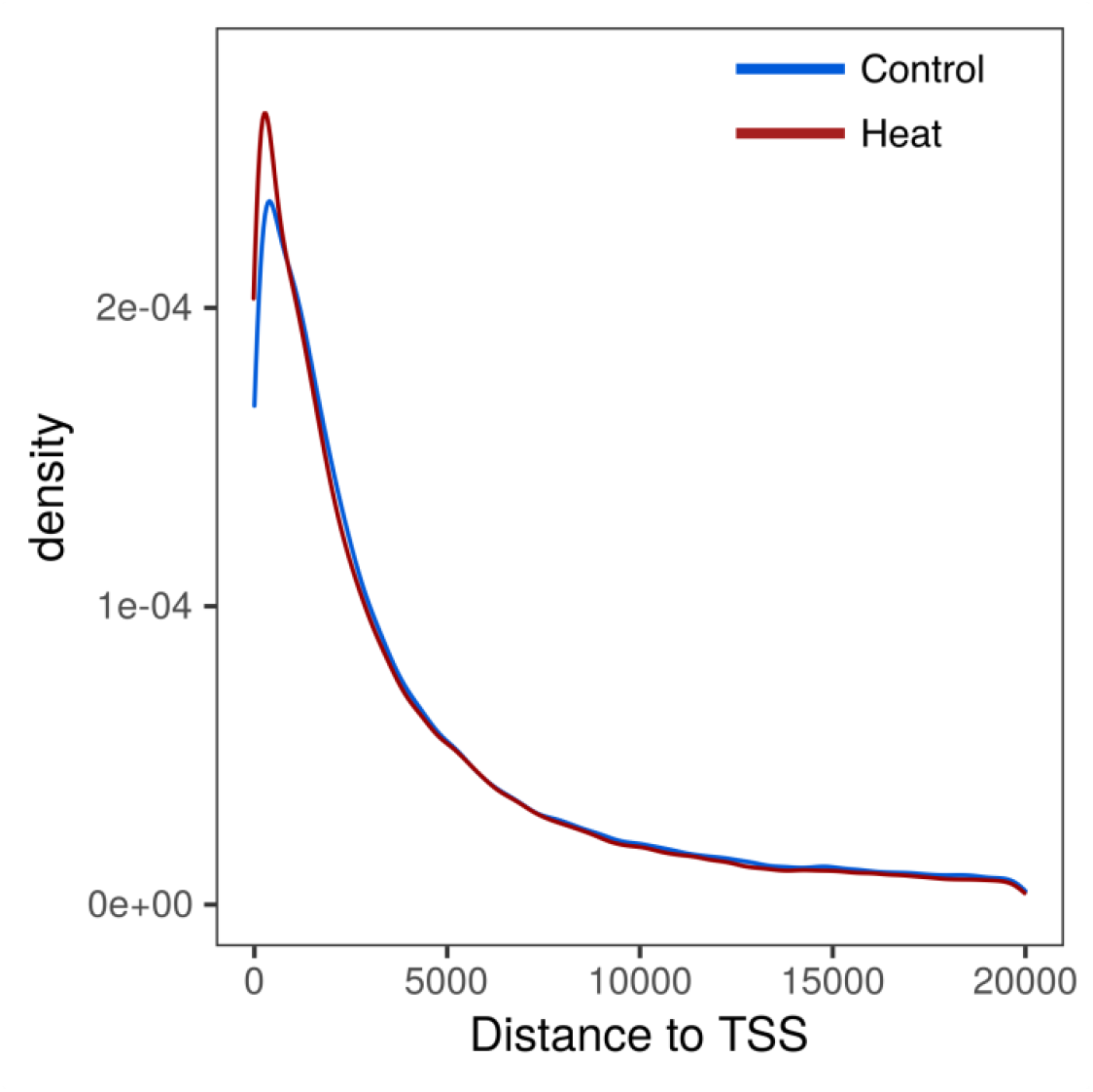
Distance of MOA-seq footprints to TSS of genes. The distance between MOA-seq TF footprint to TSS was limited to 20kb.

**Figure S9.**
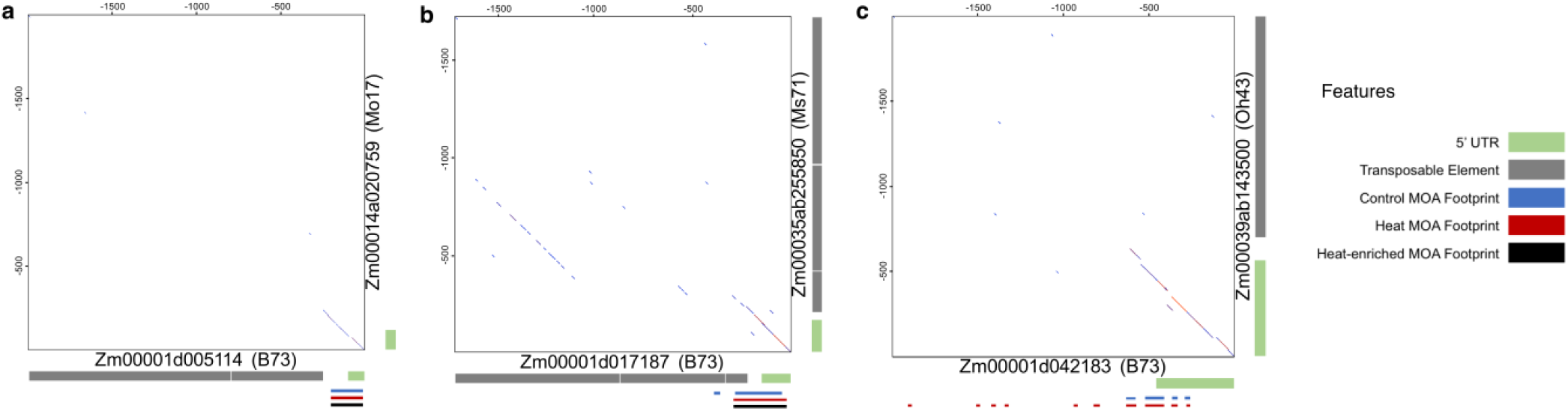
Visualization of pairwise alignments between putative promoters in pairs of reGenes. For each pair of promoter sequences that were assessed using the dual luciferase assay in protoplast systems we aligned the sequences of the two promoters. (a) *Zm00001d005114* and *Zm00014a020759;* (b) *Zm00001d017187* and *Zm00035ab255850;* (c) *Zm00001d042183 and Zm00039ab143500*. The dot plot shows the regions that are alignable as well as small insertions and deletions within these regions. The annotations of transposable elements from B73v5 and the other NAM genomes were used to document transposable elements.

### Supplemental Tables

**<u>Supplemental Table S1.</u>** BLUPs of measured photosynthetic parameters under control and heat conditions.

**<u>Supplemental Table S2.</u>** Identified genome-wide association loci using three collected photosynthetic parameters under control and heat.

**<u>Supplemental Table S3.</u>** The alignment statistics of RNA-seq samples in the large diversity panel.

**<u>Supplemental Table S4.</u>** GO enrichment for identified upregulated genes using B73 RNA-seq data collected from the large panel experiment.

**<u>Supplemental Table S5.</u>** Heat expressed only (heo) eGenes identified in the study.

**<u>Supplemental Table S6.</u>** Identified reQTLs and regulated reGenes in the study.

**<u>Supplemental Table S7.</u>** MOAseq statistics and numbers of called peaks.

**<u>Supplemental Table S8.</u>** Upregulated transposable elements overlapped with heat-enriched MOA-seq footprints in B73.

**<u>Supplemental Table S9.</u>** Enriched motifs within heat-enriched MOA-seq footprints in B73.

**<u>Supplemental Table S10.</u>** Proportion of reGenes and heo-eGenes with heat-enriched MOA-seq footprints in their 2kb flanking regions.

**<u>Supplemental Table S11.</u>** Comparison of interaction term significance between inside and outside of MOA-seq footprints in different reGene categories.

**<u>Supplemental Data 1.</u>** MOA-seq TF footprints of B73 detected in either control or heat conditions.

## Data Availability

Raw read RNA-seq data of heat stress panel have been deposited in the NCBI SRA database under the Bioproject ID PRJNA831425. MOA-seq data and associated RNA-seq data have been deposited in the NCBI SRA database under the Bioproject ID PRJNA849202. The UCSC genome browser: https://genome.ucsc.edu/s/shanwai1234/B73v4MOAseq was set up for visualizing MOA-seq and associated RNA-seq coverage of B73 samples in B73 AGPv4 genome coordinates.

## Code Availability

Codes used in this study was deposited in the Github repository: https://github.com/shanwai1234/MaizeHeatStress

## Acknowledgements

We thank Peter Hermanson, Erika Magnusson, Chase Dickson and Nicole Melamed for assistance with planting, phenotype measurements and sample collections. We thank Candice Hirsch for providing a subset of seeds from the Wisconsin Diversity Panel for the experiment and Matthew Zinselmeier for kindly providing plasmids used in Dual Luciferase reporter assays. We thank the Minnesota Agricultural Experiment Station for providing funding for the chlorophyll fluorescence camera. We thank the Minnesota Supercomputing Institute at the University of Minnesota (http://www.msi.umn.edu) for providing resources that contributed to the research results reported within this article. This study was supported by funding from the National Science Foundation (IOS-1733633 to NMS), USDA NIFA (2021-67034-35177 to ZAM), and the German Science Foundation (DFG HA 9073 to TH).

## Author contributions

Z.L. and N.M.S. conceived and designed the experiments. Z.A.M. performed the DUAL-Luciferase assay experiment, Z.L. and D.P. collected phenotype data. J.E. and T.H. performed MOA-seq library constructions and MOA-seq data processings, Z.L., Z.A.M., D.P., J.E. and T.H. analyzed data, Z.L., Z.A.M., D.P. and N.M.S. wrote the article.

